# A benchmarking framework for single-cell genome-scale metabolic model construction

**DOI:** 10.64898/2026.08.02.742346

**Authors:** Jingyu Yang, Yulong Deng, Jiahao Luo, Yonghong Wang, Feiran Li, Yu Chen

## Abstract

Single-cell genome-scale metabolic models (scGEMs) enable characterization of metabolic heterogeneity underlying cellular states and phenotypes. However, methodological choices during scGEM construction can substantially alter model structure and predictions, challenging the reliability and comparability of resulting analyses. Here, we established a systematic benchmark to assess three key construction factors: data preprocessing method, model extraction method (MEM) and gene expression threshold. We evaluated 26 strategies representing different combinations of these factors across nine scRNA-seq datasets in three dimensions: accuracy, sensitivity to expression perturbation and computational feasibility. We found that MEM had the greatest influence on most accuracy metrics, data preprocessing strongly influenced the discrimination of cellular identities, and expression threshold balanced model completeness and cellular specificity. These findings indicate that strategy performance varies across evaluation criteria. Our benchmark provides practical guidance for strategy selection and an empirical basis for standardized evaluation and future scGEM method development.

## Introduction

Metabolism is critical to diverse cellular processes, shaping cellular function, state and phenotype^1^. Different cell types and states often exhibit distinct metabolic programs, giving rise to metabolic heterogeneity within complex tissues^2^. Characterizing this heterogeneity at single-cell resolution can reveal cell-specific functional states and phenotypic differences that are obscured by bulk measurements^3^. Experimental approaches, including single-cell metabolomics and stable isotope tracing, provide increasingly powerful means to investigate cellular metabolism. However, their limited molecular coverage, high cost and restricted scalability continue to hinder systematic and quantitative characterization of metabolic network function at single-cell resolution^4,5^.

Genome-scale metabolic models (GEMs) provide a mechanistic framework for representing and simulating cellular metabolism by linking metabolites, biochemical reactions, stoichiometric constraints and gene-reaction associations^6^. Advances in single-cell RNA sequencing (scRNA-seq) have enabled the construction of single-cell GEMs (scGEMs) by integrating cell-resolved gene expression profiles with these networks. scGEMs provide cell-specific network contexts that complement gene expression profiles, enabling the assessment of metabolic functions and characterization of metabolic heterogeneity across cell types and states^7–12^. As artificial intelligence virtual cell (AIVC) frameworks increasingly seek to model cellular states and predict responses to genetic, chemical and environmental perturbations^13,14^ scGEMs may offer an interpretable metabolic layer linking predicted transcriptional states to metabolic phenotypes.

The construction of scGEMs involves several methodological factors that may affect model performance. First, scRNA-seq data are typically preprocessed before downstream analysis. Normalization is commonly used to account for variation in sequencing depth and other technical effects, whereas imputation seeks to estimate expression values unobserved owing to dropout related sparsity^15^. Both procedures alter the expression profiles used for model construction and consequently influence model performance^16^. Second, expression profiles are integrated with a reference GEM through a model extraction method (MEM) to derive context-specific models^17^. Because MEMs differ in their algorithmic assumptions and optimization objectives, they may generate networks with distinct structures and functional capacities^18^. Third, for MEMs that require expression classification, gene expression thresholds determine how genes are assigned to expression states. Genes above the threshold generally support retention of their associated reactions, whereas genes below the threshold provide weaker support and may contribute to reaction exclusion. Thresholds that are too low may retain noisy signals, whereas those that are too high may exclude biologically relevant reactions^17,18^.

The construction of reliable scGEMs remains challenging^19^. Existing studies often choose construction strategies subjectively^20–25^, which may introduce methodological variation in model structure and functional predictions. Such variation can confound biological interpretation, compromise the reliable characterization of metabolic heterogeneity and hinder comparisons across studies. These challenges are compounded by the absence of a unified gold standard for scGEM evaluation, which makes it difficult to assess construction strategies using a single definitive criterion. Therefore, a comprehensive benchmarking framework spanning multiple evaluation dimensions is needed to determine how construction strategies affect model structure, function and cell specificity, thereby providing an evidence base for robust and reproducible scGEM construction.

To address these challenges, we established a systematic benchmarking framework for scGEM construction across three key methodological factors: data preprocessing method, MEM and gene expression threshold. We evaluated 26 strategies representing different combinations of these factors across nine scRNA-seq datasets with defined biological labels or external validation data. The performance of scGEMs was assessed across three dimensions: accuracy, sensitivity to expression perturbation and computational feasibility. This benchmark provides objective-specific guidance for strategy selection and establishes a basis for more reliable and comparable scGEM analyses, standardized evaluation and future method development.

## Results

### A systematic benchmarking framework enables comprehensive evaluation of scGEM performance

To determine how construction strategies affect scGEM performance, we established a systematic benchmarking framework that evaluates combinations of construction factors using complementary performance dimensions across multiple datasets (**Fig. 1a**). The framework examined three key factors in scGEM construction: data preprocessing method, MEM and gene expression threshold. Different combinations of these factors defined 26 construction strategies, which were evaluated across nine scRNA-seq datasets with well characterized cell identities or independent validation evidence. We compared two data preprocessing methods, including the raw expression matrix (R) and the Linnorm^26^ normalized plus SAVER^27^ imputed expression matrix (LS); four MEMs, including ftINIT^11^, GIMME^28^, iMAT^29^ and rFASTCORMICS^30^; and four global percentile thresholds, including the 25th, 50th, 75th and 90th percentiles. The performance of scGEM was evaluated across three dimensions: accuracy, sensitivity to expression perturbation and computational feasibility.

**Fig. 1.**
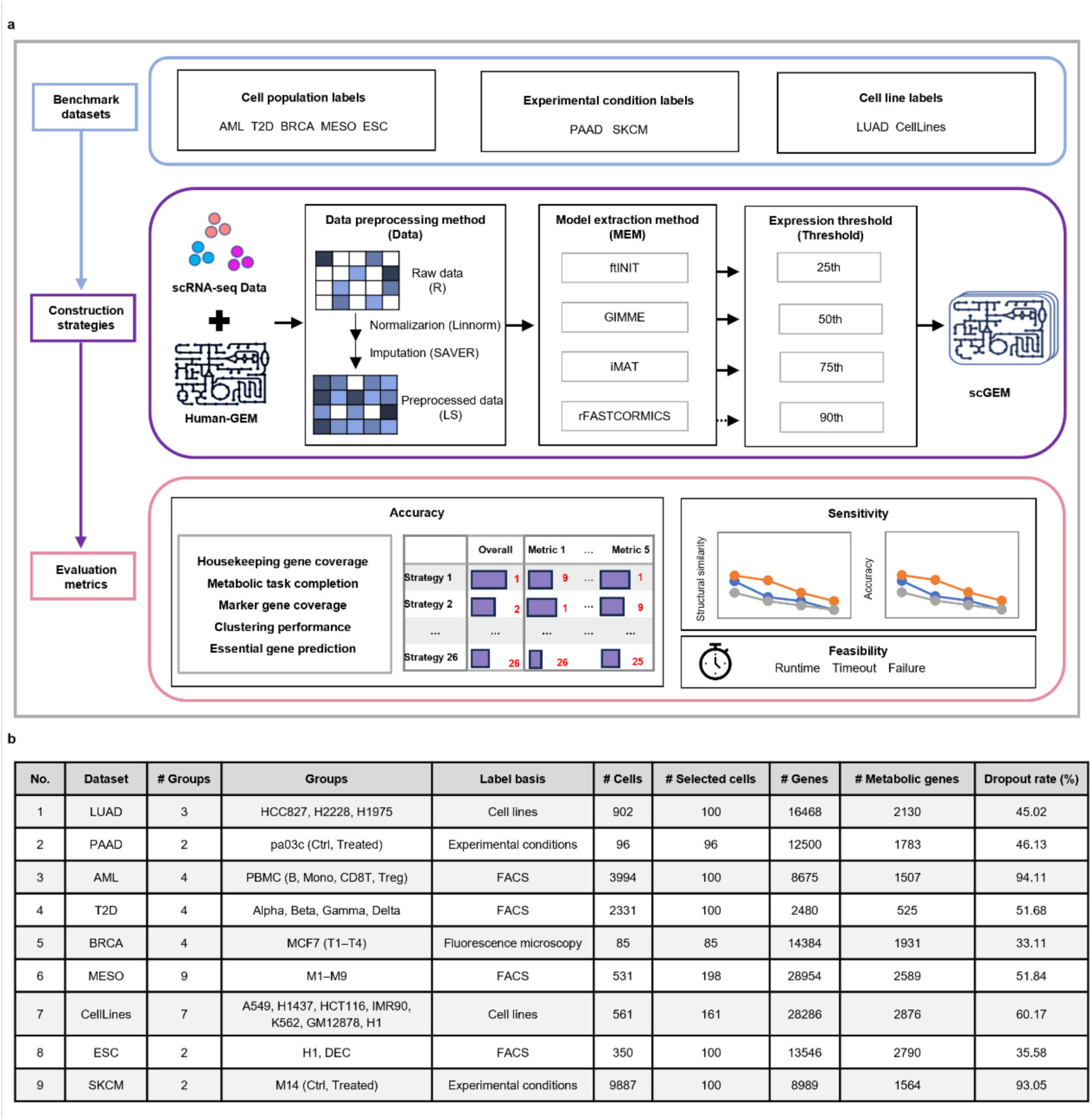
Benchmark design and scRNA-seq datasets used to evaluate scGEM construction strategies. **a**, Overview of the workflow used to benchmark scGEM construction strategies. Nine scRNA-seq datasets with established labels for cell populations, experimental conditions or cell lines were included. scGEMs were constructed by integrating scRNA-seq expression matrices with Human-GEM. 26 construction strategies were generated by varying the data preprocessing method (Data), model extraction method (MEM) and gene expression threshold (Threshold). Raw expression matrices (R) were compared with matrices normalized using Linnorm and imputed using SAVER (LS). Four MEMs were evaluated: ftINIT, GIMME, iMAT and rFASTCORMICS. Gene expression thresholds corresponding to the 25th, 50th, 75th and 90th thresholds were applied to ftINIT, GIMME and iMAT, whereas rFASTCORMICS does not require a predefined gene expression threshold. scGEM performance was assessed across three dimensions. Accuracy was evaluated using housekeeping gene coverage, metabolic task completion, marker gene coverage, clustering performance and essential gene prediction, and the resulting metric-specific scores were integrated to rank the construction strategies. Sensitivity to expression perturbation was evaluated by quantifying changes in model structure and accuracy. Computational feasibility was assessed on the basis of runtime, timeout occurrence and model construction failure. **b**, Detailed information of the nine scRNA-seq datasets included in the benchmark. LUAD, lung adenocarcinoma; PAAD, pancreatic ductal adenocarcinoma; AML, acute myeloid leukemia; T2D, type 2 diabetes; BRCA, breast cancer; MESO, mesodermal cells; CellLines, mixed cell lines; ESC, embryonic stem cells; SKCM, cutaneous melanoma. Ctrl, control; FACS, fluorescence-activated cell sorting. T1–T4 denote MCF7 cells collected at 0, 3, 6 and 12 h after 17β-estradiol stimulation, respectively. M1–M9 denote nine FACS-defined cell populations identified during mesodermal differentiation.

The construction factors were selected on the basis of their relevance to current scGEM construction workflows and their potential influence on model outputs. Previous studies have shown that normalization and imputation methods perform differently across downstream tasks, and that no single preprocessing method is consistently optimal^31,32^. Therefore, based on the study of preprocessing methods across multiple scRNA-seq analysis tasks guided by Tian et al.^31^, we selected Linnorm normalization combined with SAVER imputation as a representative preprocessing setting, and evaluated its effect on scGEM construction relative to raw expression data. MEM selection considered algorithmic principles, methodological representativeness and suitability for large scale scGEM construction. Among 14 candidate MEMs, ftINIT, GIMME, iMAT and rFASTCORMICS were selected for benchmarking, whereas the remaining methods were excluded for specific reasons (**Supplementary Note 1**). For MEMs requiring predefined expression thresholds, we evaluated four global percentile thresholds, including the 25th, 50th, 75th and 90th percentiles, to assess how expression classification stringency affects model construction. Lower thresholds classify a larger proportion of genes as expressed, whereas higher thresholds impose more stringent criteria. Because rFASTCORMICS does not require a manually specified threshold, it was excluded from threshold specific comparisons.

Evaluation metrics were organized into three complementary dimensions of scGEM performance. Accuracy was assessed using five metrics: housekeeping gene coverage, metabolic task completion, marker gene coverage, clustering performance and essential gene prediction. These metrics evaluate the retention of basal metabolic genes and core metabolic functions, the representation of cell-type-specific metabolic features, the preservation of known cellular identities, and the prediction of functional consequences following gene deletion. Sensitivity analysis examined how perturbations to input gene expression affected model structure and accuracy metrics, whereas feasibility analysis assessed construction time and success rate.

The benchmark included nine scRNA-seq datasets spanning diverse biological contexts (**Fig. 1b**). Datasets were selected on the basis of established cell-identity labels or available experimental reference data, collectively supporting evaluation across the five accuracy metrics (**Supplementary Table 1**). We prioritized annotations supported by prior biological knowledge or independent measurements rather than labels derived solely from a specific clustering or integration algorithm. The datasets covered different tissue origins, disease states, cellular compositions and dropout rates, enabling evaluation of scGEM construction strategies across diverse single cell contexts.

### Accuracy metrics are differentially influenced by scGEM construction strategies

We first evaluated the 26 construction strategies across five accuracy metrics and summarized their overall performance using normalized scores (**Fig. 2a**). Strategies were denoted in the format MEM-Data-Threshold, with each name specifying the model extraction method, data preprocessing method and gene expression threshold. Overall performance varied substantially across strategies, with ftINIT-LS-25th ranking first, followed by GIMME-LS-25th, ftINIT-LS-50th and GIMME-LS-50th. However, performance varied across individual accuracy metrics, indicating that the relative advantage of each strategy was metric dependent. Consequently, the highest overall rankings reflected balanced performance across metrics rather than superiority in any single metric.

**Fig. 2.**
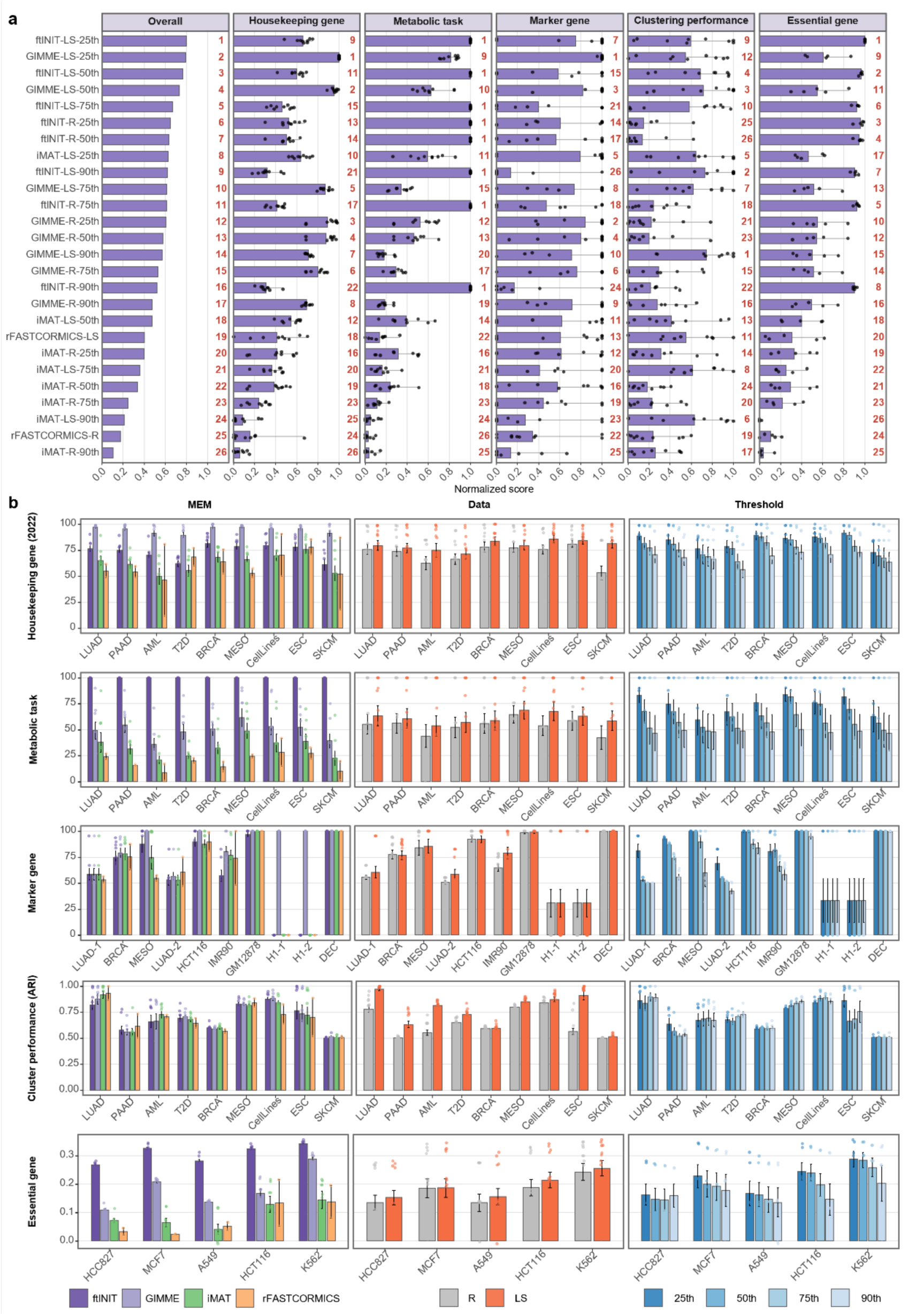
Benchmarking results for the accuracy of scGEM construction strategies. **a**, Normalized accuracy scores and overall ranking of 26 construction strategies for scGEMs across five metrics. Strategy names follow the format MEM-Data-Threshold for methods requiring a predefined gene expression threshold, where MEM, Data and Threshold denote model extraction method, data preprocessing method and gene expression threshold, respectively. rFASTCORMICS strategies follow the format MEM-Data because this method does not require an externally specified threshold. R denotes the raw expression matrix, and LS denotes the matrix normalized using Linnorm and imputed using SAVER. For each metric, raw scores were min-max normalized across strategies separately within each dataset and then averaged across datasets. Housekeeping gene coverage integrates results obtained using the gene sets published in 2019, 2022 and 2025, whereas clustering performance integrates ARI and NMI. In the five metric-specific panels, bars show the mean normalized scores across datasets, points show the normalized scores for individual datasets, and horizontal lines indicate the observed ranges across datasets. The overall score was calculated as the unweighted mean of the five metric-level scores. Red numbers indicate the rank of each strategy for the corresponding metric or overall score, with higher scores indicating better performance. The number of datasets included in each analysis depended on the availability of metric-specific reference information. Housekeeping gene coverage, metabolic task completion and clustering performance were evaluated in nine datasets; marker gene coverage in ten dataset-specific groups, with distinct cell populations or cell lines evaluated separately; and essential gene prediction in five cell lines. **b**, Raw accuracy scores grouped by construction factor across datasets. Rows correspond to the five accuracy metrics, and columns compare model extraction methods (MEM), data preprocessing methods (Data) and gene expression thresholds (Threshold). Bars show the mean raw score across strategies within each factor level and dataset, error bars indicate the standard error of the mean, and points show raw scores for individual strategies. Only datasets containing the reference information required for the corresponding metric are shown. Housekeeping gene coverage is shown using the housekeeping gene set published in 2022, and clustering performance is represented by the adjusted Rand index (ARI). Threshold comparisons include only ftINIT, GIMME and iMAT. For marker gene coverage, cell populations or cell lines within the same source dataset were treated as separate dataset-specific groups, and reference marker gene sets were matched to their biological origins. LUAD-1 and LUAD-2 correspond to the LUAD dataset and the A549 and H1437 cell lines within the CellLines dataset, respectively, whereas H1-1 and H1-2 correspond to H1 cells within the CellLines and ESC datasets, respectively.

To determine the sources of these differences, we next compared the effects of the three construction factors using the raw scores for each dataset (**Fig. 2b**, **Supplementary Fig. 1**). Data preprocessing showed the most consistent effect across the five accuracy metrics, with LS-preprocessed data generally outperforming raw expression data. This improvement was most pronounced for clustering performance (**Fig. 2b**, **Supplementary Fig. 2**). Consistently, variance decomposition identified data preprocessing as the largest contributor among the three construction factors to variation in clustering performance (**Supplementary Fig. 3**). These results indicate that preprocessing is particularly important for retaining cell-type-associated variation in scGEMs.

The choice of MEM produced more pronounced but metric-specific differences (**Fig. 2b**, **Supplementary Fig. 2**). GIMME-based strategies generally achieved higher housekeeping and marker gene coverage, whereas ftINIT-based strategies performed better for metabolic task completion and essential gene prediction. These patterns were consistent with their algorithmic designs. ftINIT explicitly incorporates predefined metabolic tasks during model extraction, which may contribute to its stronger functional performance. GIMME adopts a relatively inclusive pruning strategy, generally producing larger models with broader gene coverage. Among the three construction factors, MEM explained the largest proportion of variation in housekeeping and marker gene coverage, metabolic task completion and essential gene prediction (**Supplementary Fig. 3**). Thus, MEM choice strongly influences gene retention and metabolic functionality in scGEMs.

Gene expression threshold had a smaller but metric-specific effect (**Fig. 2b**, **Supplementary Fig. 2**). Lower thresholds, particularly the 25th percentile, generally improved housekeeping and marker gene coverage, metabolic task completion and essential gene prediction, which may reflect the contribution of lowly expressed genes to retaining network components and metabolic functional capacity. In contrast, higher thresholds tended to improve clustering performance, although the optimal threshold varied across datasets. This pattern suggests that stringent expression criteria may improve the separation of cellular identities by emphasizing more strongly expressed genes. Variance decomposition further showed that the threshold explained less variation than MEM or data preprocessing for most accuracy metrics (**Supplementary Fig. 3**). Thus, gene expression threshold primarily modulates the balance between model completeness and cellular specificity.

Together, the three construction factors influenced distinct aspects of scGEM accuracy. Data preprocessing most consistently affected the preservation of cellular identity, MEM primarily determined gene retention and metabolic functionality, and expression threshold regulated the balance between model completeness and cellular specificity. These findings demonstrate that strategy performance depends on the evaluation criteria.

### Sensitivity to expression perturbation and computational feasibility are strategy-dependent

We performed an expression perturbation sensitivity analysis to evaluate how construction strategies respond to changes in input expression profiles (**Fig. 3a**, **Supplementary Fig. 4**). An appropriate degree of sensitivity is required for a strategy to distinguish meaningful expression information from artificially perturbed inputs. More sensitive strategies produce models that more strongly reflect expression variation but may also be more affected by technical noise or preprocessing differences. Less sensitive strategies produce more structurally and functionally stable models but may incorporate less cell-specific expression information. Perturbed expression matrices were generated by progressively reducing their correlation with the original matrix. Models constructed from the perturbed data were compared with those derived from the unperturbed data in terms of structural similarity and accuracy.

**Fig. 3.**
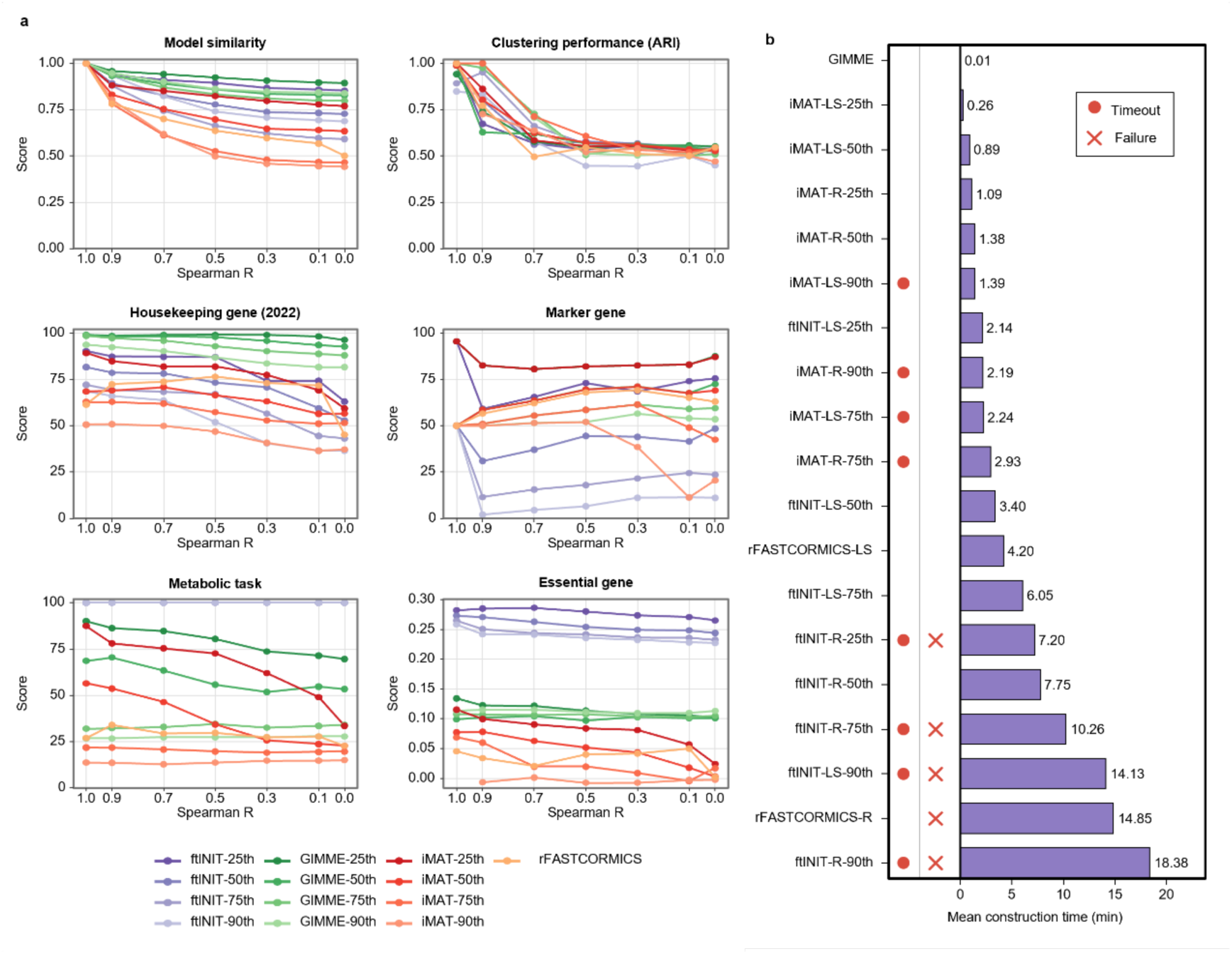
Benchmarking results for sensitivity and computational feasibility of scGEM construction strategies. **a**, Sensitivity of LS-based scGEM construction strategies to progressive expression perturbation in the LUAD dataset. The LUAD dataset was selected because it supported evaluation using all five accuracy metrics and contained biologically distinct cell-line populations. LS denotes expression matrices normalized using Linnorm and imputed using SAVER. The unperturbed expression matrix (R = 1) served as the reference. Perturbed expression matrices were generated at target Spearman rank correlations (R = 0.9, 0.7, 0.5, 0.3, 0.1 and 0) with the original matrix. Model structural similarity was quantified using the Jaccard index between the reaction sets of models constructed from perturbed expression matrices and the corresponding models constructed from the unperturbed matrix. The other panels show the corresponding raw scores for five accuracy metrics at each perturbation level. Clustering performance is represented by the adjusted Rand index (ARI), and housekeeping gene coverage is shown using the housekeeping gene set published in 2022. **b**, Computational feasibility of representative construction strategies across the benchmark datasets. Bars show the mean model construction time across datasets, and values adjacent to the bars indicate the corresponding time in minutes. Red filled circles indicate that at least one cell-specific reconstruction reached the method-specific runtime limit in any dataset, whereas red crosses indicate that at least one reconstruction returned an empty model. For ftINIT, reaching the runtime limit resulted in model construction failure and an empty model output. By contrast, iMAT could return a model upon reaching the runtime limit, and these models were retained for subsequent analyses. rFASTCORMICS failures were identified by empty model outputs without a confirmed timeout event. Empty models were excluded from the calculation of mean construction time. GIMME-based strategies had nearly identical construction times and are therefore represented by a single GIMME entry.

At the structural level, model similarity progressively decreased with increasing expression perturbation for all construction strategies, although the magnitude of change varied substantially. GIMME-based strategies retained higher structural similarity, potentially because their broader reaction retention buffered perturbation-induced changes. ftINIT-based strategies showed an intermediate response, consistent with predefined metabolic task constraints partially limiting structural changes driven by expression variation. By comparison, iMAT- and rFASTCORMICS-based strategies exhibited larger structural changes, suggesting stronger dependence on expression-based reaction classification or network consistency constraints.

At the accuracy level, perturbation effects were strongly dependent on both the metric and construction strategy. Clustering performance showed the most consistent and pronounced decline for all strategies, indicating that the retention of cellular identity variation depended strongly on the input expression patterns. Housekeeping gene coverage and essential gene prediction tended to decrease with increasing perturbation for all strategies, although the magnitude differed across MEMs and thresholds. Metabolic task completion declined more clearly with increasing expression perturbation in several GIMME- and iMAT-based strategies, whereas it showed limited sensitivity in ftINIT-based strategies. Marker gene coverage showed heterogeneous and occasionally nonmonotonic responses, which could be partly attributed to small size of the mapped marker gene sets, as the retention or loss of individual genes produced disproportionately large changes in coverage.

Overall, all construction strategies responded to expression perturbation, indicating their ability to distinguish meaningful expression information from artificially perturbed inputs. However, the degree of sensitivity varied across metrics and strategies. Structural similarity and clustering performance showed the most pronounced responses. Among the construction strategies, several iMAT-based strategies and rFASTCORMICS were relatively more sensitive, whereas GIMME- and ftINIT-based strategies generally showed smaller changes.

To assess the feasibility of large-scale scGEM generation, we compared mean construction time and timeout or failure events across strategies (**Fig. 3b**).

Computational requirements differed markedly among MEMs. GIMME was the fastest method and showed no timeout or failure events. iMAT generally required short runtimes but experienced timeouts at higher thresholds. ftINIT was slower, with computational time generally increasing under higher thresholds, and several strategies showed timeout or failure events. rFASTCORMICS was also relatively slow, particularly when applied to raw expression data. Thus, GIMME showed the most favorable computational feasibility, whereas the feasibility of the other MEMs depended more strongly on construction strategies.

### Dataset context shapes clustering performance

We next assessed the consistency of performance of strategies across datasets. Strategy rankings were highly concordant for metabolic task completion, housekeeping gene coverage and essential gene prediction, but substantially less concordant for marker gene coverage and clustering performance (**Fig. 4a**). Pairwise rank correlations for the latter two metrics also varied widely across dataset pairs (**Fig. 4b**, **Supplementary Fig. 5**), indicating that metrics reflecting cell-specific features were more dependent on dataset context. The relatively low concordance in marker gene coverage may partly reflect the small size of several dataset-specific marker gene sets. Because many sets contained only a few mapped genes, coverage scores were discrete and many strategies received identical scores (**Supplementary Fig. 6**), such that the retention or loss of only a small number of genes could substantially alter strategy rankings.

**Fig. 4.**
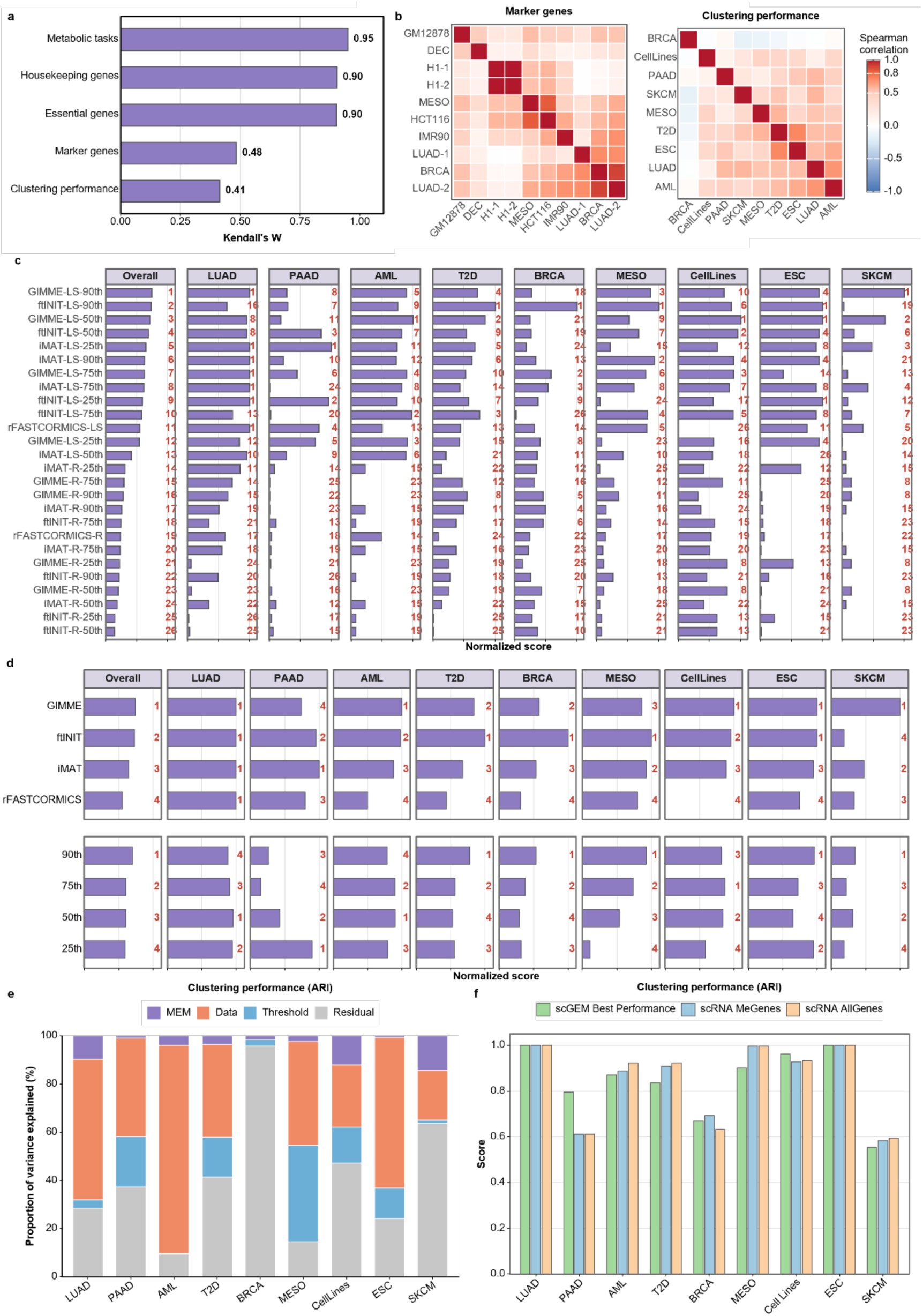
Dataset-dependent performance of scGEM construction strategies. **a**, Cross-dataset concordance of construction-strategy rankings across five accuracy metrics. Concordance was quantified using Kendall’s coefficient of concordance (W). Values approaching 1 indicate greater consistency in strategy rankings across datasets, whereas values approaching 0 indicate stronger dataset dependence. **b**, Pairwise similarity of construction-strategy performance across datasets for marker gene coverage and clustering performance. These two metrics showed the lowest cross-dataset concordance in **a**. Each heatmap cell shows the Spearman rank correlation between the scores of the 26 construction strategies for a pair of datasets. **c**, Normalized clustering scores and rankings of the 26 construction strategies across nine datasets. Adjusted Rand index (ARI) and normalized mutual information (NMI) values were separately min-max normalized across strategies within each dataset and then averaged to obtain an integrated clustering score. The Overall panel shows the mean integrated score across datasets, whereas the other panels show dataset-specific integrated scores. Bars show integrated clustering scores, and red numbers indicate the rank of each strategy within the corresponding panel, with higher scores indicating better performance. Strategy names follow the format MEM-Data-Threshold for methods requiring a predefined gene expression threshold, where MEM, Data and Threshold denote the model extraction method, data preprocessing method and gene expression threshold, respectively. rFASTCORMICS strategies follow the format MEM-Data because this method does not require a predefined gene expression threshold. R denotes the raw expression matrix, and LS denotes the matrix normalized using Linnorm and imputed using SAVER. **d**, Factor-level comparison of integrated clustering scores using LS-preprocessed expression matrices. In the MEM comparison, each threshold-dependent method was represented by its highest-scoring threshold setting within each dataset. In the Threshold comparison, scores were averaged across ftINIT, GIMME and iMAT; rFASTCORMICS was excluded because it does not require an externally specified threshold. The Overall panels show mean scores across datasets, and the remaining panels show dataset-specific scores. Bars show normalized clustering scores, and red numbers indicate the rank of each factor level within the corresponding panel. **e**, Variance decomposition of raw ARI values across benchmark datasets. Stacked bars show the proportion of variance explained by MEM, Data and Threshold, with unexplained variation represented by the residual. rFASTCORMICS was excluded from analyses involving Threshold. **f**, Comparison of scGEM-based and expression-based clustering performance. For each dataset, the scGEM result corresponds to the construction strategy with the highest integrated clustering score in **c**. Expression-based clustering was performed using scRNA-seq expression profiles restricted to metabolic genes represented in Human-GEM or using all detected genes.

We therefore examined clustering performance, which showed the lowest cross-dataset concordance, separately for each dataset. Strategy rankings varied markedly, and the leading strategy varied across all datasets (**Fig. 4c**). Because LS-based strategies generally outperformed those based on raw expression data (**Supplementary Fig. 7**), subsequent comparisons of MEM and threshold effects were restricted to LS- preprocessed data. The best-performing MEM differed across datasets (**Fig. 4d**), demonstrating that the relative performance of MEM was context dependent. Higher expression thresholds generally improved clustering, although lower or intermediate thresholds were favored in several datasets. These results suggest that the optimal combination of MEM and threshold depends on the biological and technical characteristics of the input dataset. Variance decomposition further showed that the relative contributions of data preprocessing, MEM and threshold varied substantially across datasets (**Fig. 4e**, **Supplementary Fig. 8**). Thus, clustering performance was shaped by dataset-specific effects of the construction factors rather than by a uniform construction rule.

Finally, to evaluate whether scGEMs retain the intrinsic cell-type separability present in the transcriptomic data, we compared the best-performing strategy for each dataset with clustering based directly on metabolic genes or all genes in the corresponding scRNA-seq data (**Fig. 4f**, **Supplementary Fig. 9**). The best-performing scGEMs achieved performance comparable to expression-based clustering in several datasets, indicating that scGEMs constructed by suitable strategies can preserve biologically relevant information about cellular identity.

### Model size influences the interpretation of scGEM accuracy metrics

Because construction strategies generated models of different sizes, we next examined whether model size influenced performance evaluation. The numbers of reactions, genes and metabolites varied markedly among strategies (**Fig. 5a**, **Supplementary Fig. 10**). GIMME generated the largest models, likely because its permissive pruning strategy tends to retain more reactions. ftINIT and iMAT generated intermediate-sized models, although through different mechanisms: ftINIT incorporates metabolic task constraints to maintain functional consistency, whereas iMAT balances the inclusion of highly expressed reactions with the exclusion of lowly expressed reactions. In contrast, rFASTCORMICS produced the smallest models, consistent with its construction of compact, consistency-based subnetworks from expression-supported core reactions. The three size measures showed similar trends and were highly positively correlated (**Supplementary Fig. 11**).

**Fig. 5.**
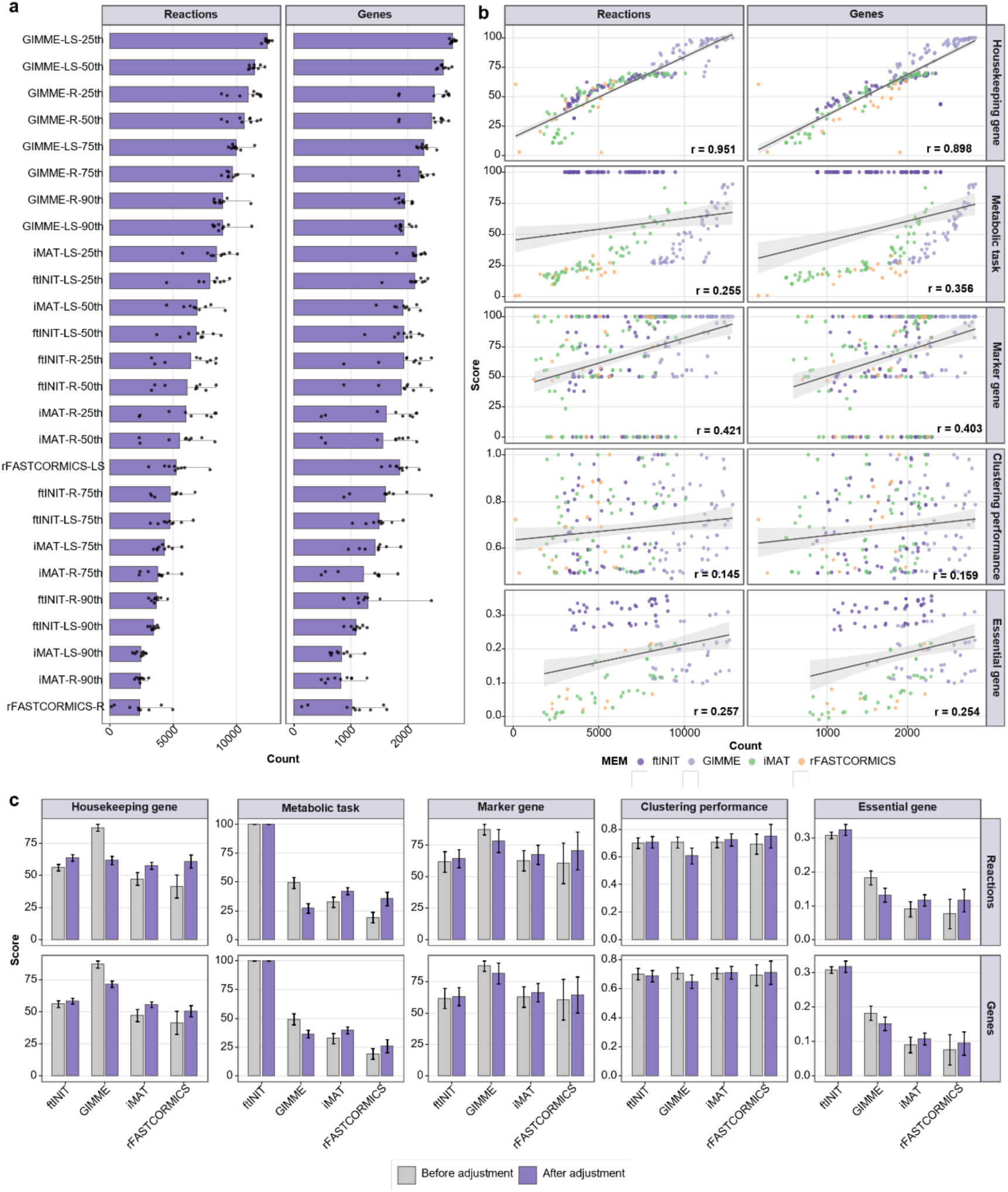
Model-size effects on scGEM accuracy metrics. **a**, Reaction and gene counts of models generated by 26 construction strategies. Strategy names follow the format MEM-Data-Threshold for methods requiring a predefined gene expression threshold, where MEM, Data and Threshold denote the model extraction method, data preprocessing method and gene expression threshold, respectively. rFASTCORMICS strategies follow the format MEM-Data because this method does not require a predefined gene expression threshold. R denotes the raw expression matrix, and LS denotes the matrix normalized using Linnorm and imputed using SAVER. For each strategy, reaction and gene counts were first averaged across cell-specific models within each dataset. Bars show the mean of these dataset-level averages across datasets, and points show values for individual datasets. Strategies are ordered by mean reaction number. **b**, Correlations between reaction or gene number and raw accuracy metrics across strategy-dataset combinations. Each point represents one construction strategy in one dataset and is colored according to MEM. Grey lines show linear regression fits, with shaded areas indicating 95% confidence intervals. Spearman correlation coefficients are shown in the corresponding panels. Housekeeping gene coverage was calculated using the housekeeping gene set published in 2022, and clustering performance was represented by the adjusted Rand index (ARI). **c**, MEM-level accuracy performance before and after adjustment for model size. Separate linear models were fitted for each accuracy metric using the raw metric score as the response and MEM and model size as explanatory variables. Reaction number and gene number were included separately as model-size covariates in the upper and lower rows, respectively. Error bars indicate 95% confidence intervals.

Accuracy metrics showed different degrees of association with model size (**Fig. 5b**, **Supplementary Fig.12**). Housekeeping gene and marker gene coverage increased with gene numbers, as larger models tended to retain more genes. Metabolic task completion and essential gene prediction were also positively associated with reaction number. These associations became more pronounced after excluding ftINIT, which frequently achieved high scores because of task-related constraints (**Supplementary Fig.13**). By comparison, clustering performance showed only weak correlation with model size, suggesting that it is less affected by model scale and may be more suitable for evaluating the preservation of cell-state differences.

To account for model size in comparisons among MEMs, we estimated size-adjusted performance using linear models (**Fig. 5c**, **Supplementary Fig.14**). GIMME performance declined in most metrics after size adjustment, indicating that its apparent advantage was partly driven by larger model size, although it remained superior for gene-coverage metrics. Conversely, iMAT and rFASTCORMICS achieved higher adjusted scores for several metrics, indicating that smaller-model methods may be underestimated by raw scores. ftINIT retained high performance in metabolic task completion and essential gene prediction, suggesting that its functional advantage was not primarily driven by model scale. Clustering performance changed only slightly after adjustment, consistent with its weak association with model scale. Together, these results demonstrate that model size can confound several scGEM accuracy metrics and should therefore be considered when comparing construction methods.

### Benchmark results guide scGEM construction and method development

Based on the systematic benchmark, we provide objective specific guidelines for scGEM construction (**Fig. 6)**. Our results support that scRNA-seq data should be appropriately preprocessed before model construction rather than using raw expression matrices directly. For applications focused on housekeeping or marker gene coverage, GIMME-LS-25th is recommended. For metabolic task completion and essential gene prediction, ftINIT-LS-25th should be prioritized. For the discrimination of cell types, preprocessing is the key consideration, with GIMME or ftINIT selected as the MEM and the expression threshold optimized for the dataset. Although the 75th- or 90th- percentile thresholds generally performed well for clustering, lower thresholds remained preferable in some datasets. When computational efficiency is the primary consideration, GIMME is recommended prioritized because of its short runtime and low incidence of timeout or failure events. Together, scGEM construction strategies should be selected according to the intended research objective rather than applied uniformly across applications.

**Fig. 6.**
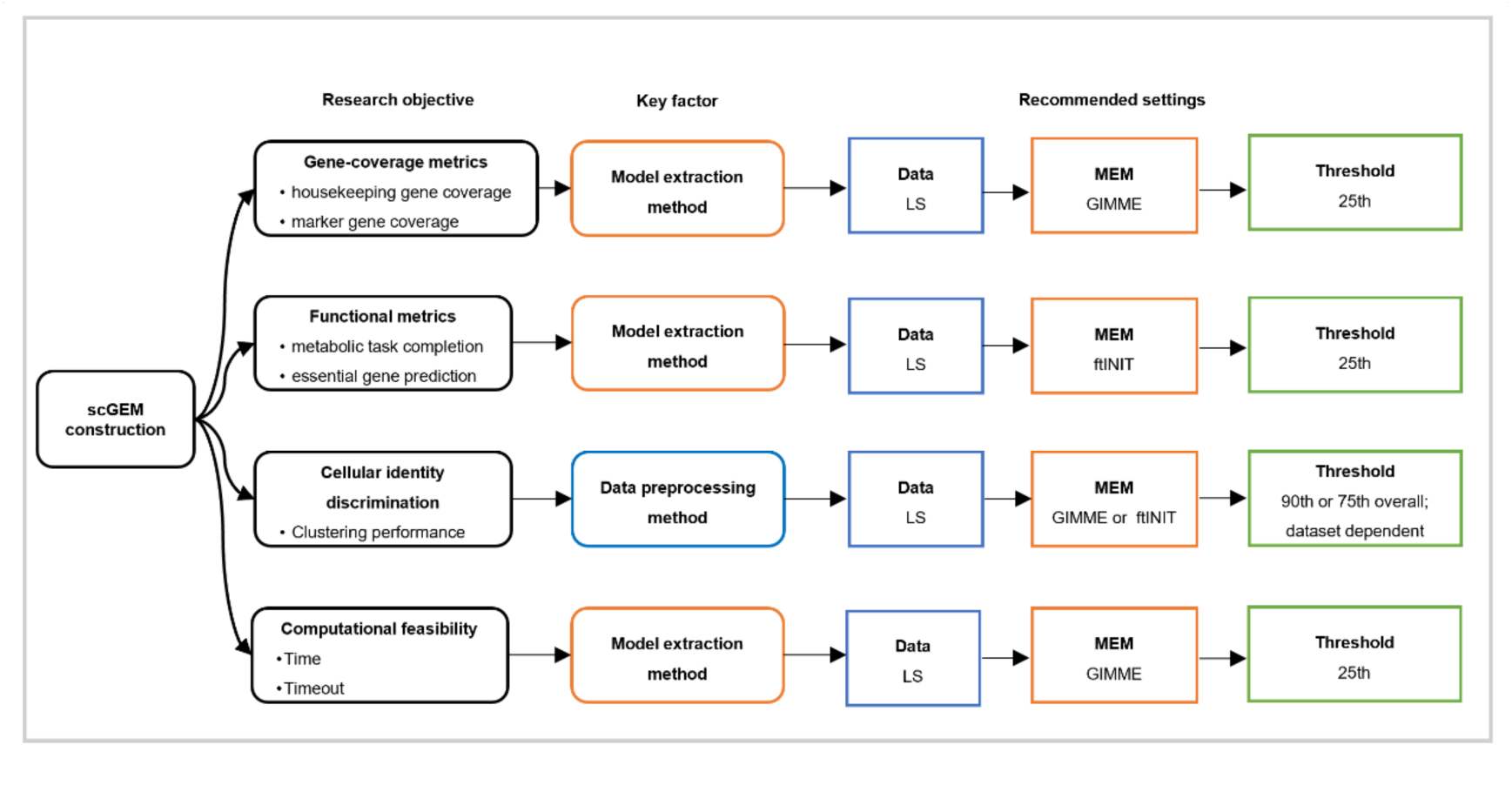
Objective-driven guidance for selecting scGEM construction strategies. Schematic summary of recommended settings for scGEM construction according to different research objectives. Across the objectives shown, scRNA-seq expression matrices should generally be preprocessed before model construction. For gene-coverage metrics, evaluated using housekeeping and marker gene coverage, MEM is the primary determinant, with GIMME-LS-25th recommended. For functional objectives, including metabolic task completion and essential gene prediction, ftINIT-LS-25th is recommended. For cellular identity discrimination based on clustering performance, data preprocessing is the key factor. LS input with either GIMME or ftINIT is recommended, and the threshold should be optimized for the dataset. Thresholds corresponding to the 75th or 90th provide reasonable initial settings, although lower thresholds may be preferable for some datasets. For computational feasibility, assessed by model construction time and timeout occurrence, GIMME is prioritized on the basis of shorter construction times and fewer timeout events. Data, data preprocessing method; MEM, model extraction method; Threshold, gene expression threshold; LS, expression matrix normalized using Linnorm and imputed using SAVER.

Beyond guiding the selection of existing construction strategies, the benchmark provides conceptual directions for future scGEM method development. Future MEMs should balance gene retention, metabolic functionality and the resolution of cell specific differences. Data preprocessing should be treated as an integral component of scGEM construction and optimized jointly with model extraction to mitigate sparsity and technical variation in scRNA seq data. Threshold selection should also move beyond fixed global cutoffs toward adaptive thresholds or continuous expression weighting schemes that account for dataset and objective dependence. Future methods may further combine complementary algorithmic features, such as broad network retention and explicit metabolic task constraints. These developments could improve the reliability, interpretability and applicability of scGEMs across diverse single cell contexts.

## Discussion

In this study, we established a systematic benchmarking framework to evaluate how key construction factors affect scGEM performance across diverse single-cell datasets. By comparing 26 strategies across nine scRNA-seq datasets, we assessed model accuracy, sensitivity to expression perturbation and computational feasibility. The three construction factors made distinct contributions to model performance: MEM choice primarily shaped gene retention and metabolic functionality, data preprocessing most strongly affected clustering performance, and gene expression threshold regulated the balance between model completeness and the discrimination of cellular identities. These findings indicate that different construction strategies are suited to different research objectives and provide an evidence base for objective-specific scGEM construction and method development.

Existing studies have largely relied on MEMs originally developed for bulk omics to construct scGEMs from scRNA-seq data^20–25^. However, these studies generally focused on specific modeling pipelines or biological applications. Recent studies further questioned the extent to which methods developed for bulk omics are applicable to single-cell data, given the additional challenges posed by sparsity, technical variation and cellular heterogeneity, and whether fundamentally different modeling approaches may be required^19,33^. By comparing alternative construction strategies across complementary evaluation dimensions, our benchmark provides systematic evidence on the applicability and limitations of existing approaches to scGEM construction.

Previous benchmarks using bulk RNA-seq data have shown that MEM choice strongly influences GEM composition and predictive performance, with different methods favored by different evaluation criteria^17,18^. Our study similarly found that MEM was the dominant methodological factor for several gene retention and functional metrics across diverse scRNA-seq datasets. In particular, bulk omics studies also identified GIMME and tINIT-based methods as among the best-performing approaches, although their relative strengths varied across evaluation dimensions^34,35^. Our benchmark revealed a similar pattern for GIMME and ftINIT, an optimized implementation of tINIT. Both methods performed well overall but showed distinct metric-specific advantages. This concordance between bulk and single-cell benchmarks suggests that the core principles of bulk-derived MEMs are transferable to scGEM construction, although their implementation requires adaptation to the statistical properties of scRNA-seq data. These results provide an empirical basis for refining existing MEMs and guiding the development of methods better suited to single-cell contexts.

Notably, some findings at single-cell resolution differed from those reported in bulk studies. A previous bulk benchmark found that more stringent expression thresholds generally improved essential gene prediction accuracy^17^. By contrast, lower thresholds achieved higher accuracy in our scGEM benchmark, suggesting that stringent exclusion of low-expression signals may discard functionally relevant information from sparse scRNA-seq data. In addition, a large-scale benchmark of bulk cancer models showed that models generated using different MEMs consistently distinguished tissue and cancer types with high accuracy^35^, indicating that these broad biological differences were robust to MEM choice. In our study, however, clustering performance varied substantially across datasets and construction strategies and was particularly influenced by data preprocessing. This finding suggests that methodological choices have a greater influence on the characterization of subtle metabolic variation at single-cell resolution than on that of broader variation in bulk data.

Data preprocessing remains underexplored systematically in scGEM construction. Existing studies often aggregate expression profiles across cells of the same type into pseudo-bulk inputs to reduce sparsity in scRNA-seq data^11,12,22,23^. Only a few studies have applied normalization or imputation before reconstructing models for individual cells^20,21,24^, and the impact of such preprocessing remains poorly understood. By comparing raw and LS-preprocessed inputs across multiple MEMs and accuracy metrics, our benchmark showed that preprocessing improved performance across all five accuracy metrics on average, with the largest improvement observed for clustering performance. This finding indicates that preprocessing is particularly consequential when scGEMs are used to distinguish metabolic differences among cell types. However, because our comparison was limited to LS and raw data, these results do not establish LS as universally optimal. Instead, our findings highlight data preprocessing as an essential methodological consideration in scGEM construction.

Recent work on AIVCs has emphasized the need to complement cellular state prediction with mechanistic explanations of the underlying biological processes^13^. Mechanistic virtual-cell frameworks have likewise been proposed to connect intracellular states and extracellular signals with biochemical processes and cellular behavior^14^. In this context, scGEMs could provide a constraint-based metabolic layer that translates AIVC-predicted transcriptomic states into interpretable metabolic network alterations and associated cellular phenotypes. Our benchmark shows that the reliability of this layer depends on data preprocessing, MEM choice and gene expression threshold, and provides objective-specific principles for evaluating and integrating metabolic components into future AIVC frameworks.

## Methods

### scRNA-seq datasets for benchmarking

We collected nine scRNA-seq datasets with established cell identities or independent validation data to evaluate the performance of different scGEM construction strategies. All datasets were obtained from public repositories, including the NCBI Gene Expression Omnibus (GEO) and Figshare.

Specifically, the LUAD dataset contains three lung adenocarcinoma cell lines, including HCC827, H2228 and H1975, and was obtained from GEO under accession GSE118767. The PAAD dataset was derived from pancreatic ductal adenocarcinoma samples and includes two experimental conditions, control and APEX1-KD, and was obtained from GEO under accession GSE99305. The AML dataset was derived from peripheral blood mononuclear cells of patients with acute myeloid leukemia and includes four cell populations: B cells, CD14 monocytes, naive cytotoxic T cells and regulatory T cells. This dataset was obtained from Figshare via https://doi.org/10.6084/m9.figshare.11787210. The T2D dataset was derived from type 2 diabetes samples and includes four pancreatic cell types, Alpha, Beta, Gamma and Delta, and was obtained from GEO under accession GSE85241. The BRCA dataset was derived from MCF7 invasive breast cancer cells stimulated with 17β-estradiol at four time points, 0, 3, 6 and 12 h, and was obtained from GEO under accession GSE107858. The MESO dataset contains nine mesodermal differentiation stages and was obtained from Figshare via https://doi.org/10.6084/m9.figshare.11787210. The CellLines dataset contains seven cell lines, including the lung adenocarcinoma cell lines A549 and H1437, the colorectal cancer cell line HCT116, the lung fibroblast cell line IMR90, the chronic myeloid leukemia cell line K562, the lymphoblastoid cell line GM12878 and the human embryonic stem cell line H1_ESC, and was obtained from GEO under accession GSE81861. The ESC dataset includes human embryonic stem cells H1_ESC and definitive endoderm cells and was obtained from GEO under accession GSE75748. The SKCM dataset was derived from the melanoma M14 cell line and includes control and vemurafenib-treated cells, and was obtained from GEO under accession GSE134838.

For each dataset, the expression matrix and corresponding cell or condition labels were retained for downstream scGEM construction and evaluation. Dataset labels were classified according to their annotation basis, including cell line labels, experimental condition labels and cell population labels. These labels were used as reference annotations for clustering evaluation, whereas additional external information was used when available for other accuracy metrics, including marker gene coverage and essential gene prediction.

### Quality control and preprocessing of scRNA-seq data

For each scRNA-seq dataset, quality control was performed at both the gene and cell levels before model construction. A gene was considered detected when its expression value was greater than 0. Genes were retained if they were detected in more than ten cells and had a mean expression value greater than 1 among cells in which they were detected. Cells with more than 100 detected genes were retained. The resulting filtered, unnormalized expression matrix was denoted R. To generate the LS matrix, the complete filtered expression matrix was first normalized using Linnorm^26^ and subsequently imputed using SAVER^27^. The resulting Linnorm-normalized and SAVER-imputed expression matrix was denoted LS.

For each dataset, a dataset-specific number of cells was selected for scGEM construction using a fixed random seed. Sampling was stratified by the reference cell-population, experimental-condition or cell-line labels, with the numbers of selected cells balanced as evenly as possible across groups. For datasets with relatively few cells, all available cells were retained. The complete R and LS matrices were generated before cell selection, and the same sampled cells were subsequently extracted from both matrices to ensure matched comparisons between R- and LS-based construction strategies.

### Construction of scGEMs

Human-GEM^36^ (version 1.16) was used as the reference GEM. All constructions were performed in MATLAB using the COBRA Toolbox^37^ and Gurobi Optimizer 13.0.1. Gene identifiers in each expression matrix were matched to those in Human-GEM, and genes not represented in Human-GEM were excluded from model extraction. For each selected cell, the corresponding expression profile was obtained from either the R or LS matrix and used to construct a cell-specific scGEM.

The threshold-dependent MEMs, i.e., ftINIT^11^, GIMME^28^ and iMAT^29^, were evaluated using two data preprocessing methods and four gene expression thresholds corresponding to the 25th, 50th, 75th and 90th percentiles. rFASTCORMICS^30^ was evaluated using the two data preprocessing methods without an externally specified gene expression threshold. These combinations yielded 26 scGEM construction strategies. Each cell-specific scGEM was constructed independently under every applicable strategy, and model construction time was recorded for each reconstruction.

For each dataset, percentile-based gene-expression cutoffs were calculated separately for the R and LS matrices using all expression values, including zero, across all retained genes and cells. The cutoffs were calculated separately for each dataset and preprocessing method but were shared across all cells within the corresponding matrix.

### ftINIT

ftINIT-based scGEMs were generated using RAVEN^38^ and a precomputed Human-GEM structure produced with prepINITModel. For each input matrix, the corresponding global gene-expression cutoff was supplied through the threshold field of the ftINIT transcriptomic input structure. Models were generated using the 1+0 configuration specified by getHumanGEMINITSteps, without proteomic or metabolomic input.

### GIMME

For each percentile-based gene-expression cutoff, expression values less than or equal to the cutoff were set to zero, whereas values above the cutoff were retained. Gene-level expression values were mapped to reaction-level scores according to gene-protein-reaction (GPR) rules, using the minimum available value for AND relationships and the maximum value for OR relationships. Reactions without mapped expression information were assigned a score of -1. The resulting scores were supplied to createTissueSpecificModel, with GIMME specified as the MEM and the reaction-level expression threshold fixed at 0.

### iMAT

Gene-expression thresholding and GPR mapping were performed as described for GIMME. The resulting reaction-level scores were supplied to createTissueSpecificModel, with iMAT specified as the MEM. The lower and upper reaction-level expression thresholds were both fixed at 1×10^-10^. Accordingly, reactions with a score of 0 were classified as lowly expressed, reactions with a positive score as highly expressed and reactions assigned -1 as lacking mapped expression information. We used an adapted binary implementation of iMAT in which reactions were classified as either lowly or highly expressed, without an intermediate-expression category. This implementation ensured that expression classification was defined consistently at the gene level across threshold-dependent MEMs.

### rFASTCORMICS

Expression values were discretized using discretize_FPKM. The discretized profiles were integrated with a flux-consistent version of Human-GEM using fastcormics_RNAseq and the rFASTCORMICS gene-mapping dictionary.

Because rFASTCORMICS applies its own discretization procedure, no external percentile-based gene-expression threshold was specified.

### Accuracy evaluation metrics

Five complementary metrics were used to evaluate scGEM performance: housekeeping gene coverage, metabolic task completion, marker gene coverage, clustering performance and essential gene prediction.

### Housekeeping gene coverage

This metric was used to assess the retention of genes associated with fundamental cellular functions. Three published single-cell housekeeping gene sets reported in 2019^39^ (https://sydneybiox.github.io/scMerge), 2022^40^ (http://61.160.194.165:3080/hSEGdb) and 2025^41^ (https://v24.proteinatlas.org/humanproteome/tissue/tissue+specific) were used as references. For each scGEM, coverage was calculated as the proportion of these reference genes retained in the model. Coverage values were averaged across scGEMs to obtain the score for each construction strategy.

### Metabolic task completion

This metric was used to assess the retention of essential metabolic functions required for cell survival. A set of 57 essential metabolic tasks provided by Human-GEM^36^ was used as the reference task set. For each scGEM, the tasks were evaluated individually, and metabolic task completion was calculated as the proportion of successfully completed tasks among all 57 reference tasks. Completion rates were averaged across cell-specific models to obtain the score for each construction strategy.

### Marker gene coverage

This metric was used to assess the retention of metabolic marker genes associated with specific cell types or states. Marker genes for eight target populations were obtained from the human entries in the CellMarker 2.0 database^42^. Each marker gene set was intersected with the genes represented in Human-GEM, and the mapped genes were retained as the reference metabolic marker set. Eight resulting metabolic marker gene sets were evaluated across ten dataset-population contexts. Specifically, lung adenocarcinoma markers were evaluated in the LUAD dataset (LUAD-1) and in A549 and H1437 cells from the CellLines dataset (LUAD-2), whereas H1 embryonic stem cell markers were evaluated in H1 cells from the CellLines (H1-1) and ESC (H1-2) datasets. For each scGEM, marker gene coverage was calculated as the proportion of reference marker genes retained in the model. Coverage values were averaged across cell-specific models to obtain the score for each construction strategy.

### Clustering performance

This metric was used to assess the preservation of differences among cell types or states. For each dataset and construction strategy, a binary reaction-presence matrix was generated from the reaction composition of the cell-specific models. Cells were clustered on the basis of this matrix, and the resulting cluster assignments were compared with the corresponding reference cell labels or experimental groups. Clustering performance was quantified using the ARI and NMI. ARI ranges from -1 to 1 and NMI from 0 to 1, with higher values indicating greater agreement between model-derived clusters and the reference annotations. ARI and NMI were calculated separately for each applicable dataset and construction strategy.

### Essential gene prediction

This metric was used to assess the ability of scGEMs to predict gene essentiality under metabolic perturbation. For each scGEM, metabolic genes were individually knocked out according to the GPR rules, and the resulting model was evaluated against the 57 essential metabolic tasks defined in Human-GEM. A gene was classified as model-predicted essential if its knockout prevented completion of at least one essential metabolic task.

Model predictions were compared with experimental CRISPR knockout data for five benchmark cell lines: HCC827, MCF7, A549, HCT116 and K562. Experimental essentiality data for HCT116 were obtained from the Hart 2015 dataset^43^, whereas data for the remaining cell lines were obtained from DepMap database^44,45^. For DepMap, genes with gene effect scores below -0.6 were classified as experimentally essential, following the criterion used in the Human-GEM study, and all remaining genes were classified as non-essential.

For each scGEM, model-predicted and experimental essentiality labels were matched to construct a confusion matrix. Prediction performance was quantified using the Matthews correlation coefficient (MCC)^46^:

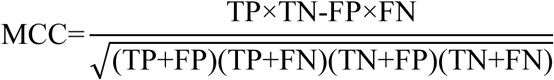

where TP, TN, FP and FN denote true positives, true negatives, false positives and false negatives, respectively. MCC values were averaged across cell-specific models to obtain the essential gene prediction score for each construction strategy. Higher MCC values indicate greater agreement between model-predicted and experimentally observed gene essentiality.

### Normalization and ranking of accuracy scores

To enable comparison of the 26 construction strategies across accuracy metrics with different numerical scales, raw scores were min-max normalized separately for each dataset and metric across construction strategies:

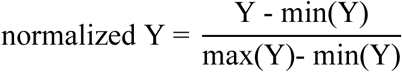

where Y denotes the raw metric score, and min(Y) and max(Y) denote the minimum and maximum raw scores across construction strategies for the same metric and dataset.

For each construction strategy and metric, normalized scores were averaged across all applicable datasets, with each dataset contributing equally, to obtain the metric-level score. The overall score was calculated as the unweighted mean of the five metric-level scores. For metrics comprising multiple submetrics, each submetric was normalized separately before equal-weight averaging. Specifically, scores derived from the three housekeeping gene sets were averaged to obtain the housekeeping gene coverage score, whereas separately normalized ARI and NMI values were averaged to obtain the clustering performance score. Normalized scores were used for cross-metric comparison, visualization and strategy ranking, whereas raw scores were retained for metric-specific analyses unless otherwise stated. Strategies were ranked in descending order of score, with tied values assigned the same rank.

### Expression perturbation sensitivity analysis

To evaluate the sensitivity of scGEM construction strategies to perturbations in input gene expression, we used the LS matrix from the LUAD dataset. This dataset was selected because it contained the reference information required for all five accuracy metrics and comprised biologically distinct cell-line populations. The matrix contained 16,468 genes and 100 cells, and perturbations were generated independently for each cell.

For each cell, the original expression vector ***x*** was randomly permuted across genes to generate a randomized vector ***r***, thereby preserving the distribution of expression values while disrupting their gene assignments. The perturbed expression vector was defined as

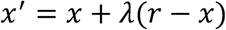

where *λ* ∈[0,1] controls perturbation strength. For each cell and target perturbation level, *λ* was selected to produce a perturbed vector whose Spearman correlation *R* with the original vector was closest to the target value. Target correlations were 1.0, 0.9, 0.7, 0.5, 0.3, 0.1 and 0, with lower values representing stronger perturbation. For *R* =1.0, the original expression vector was retained.

At each perturbation level, scGEMs were reconstructed using the same construction strategies applied to the unperturbed LS matrix. For each cell, structural similarity between the perturbed and corresponding unperturbed model was quantified using the Jaccard index of retained reaction sets. The five accuracy metrics were then recalculated to characterize changes in model performance with increasing expression perturbation.

### Computational feasibility analysis

Computational feasibility was evaluated using model construction time, timeout occurrence and model construction failure. Construction time was recorded separately for each cell-specific reconstruction. For each construction strategy and dataset, mean construction time was calculated across successfully generated cell-specific models, and the dataset-level means were subsequently averaged with equal weight to obtain the strategy-level construction time.

The default runtime limits of the corresponding implementations were 5,000 s for ftINIT and 7,200 s for iMAT. ftINIT reconstructions reaching the runtime limit were considered unsuccessful and returned empty models. By contrast, iMAT could still return a model upon reaching the runtime limit, and these models were retained for subsequent analyses. rFASTCORMICS could also fail to generate a valid model and return an empty model without a confirmed timeout event. Empty models generated by ftINIT or rFASTCORMICS were excluded from subsequent model-based analyses and from the calculation of mean construction time.

### Model-size quantification and size-adjusted performance analysis

Model size was quantified for each scGEM using the numbers of reactions, genes and metabolites. For each construction strategy and dataset, model-size measures were averaged across successfully generated cell-specific models, with empty models excluded. Associations between model size and each raw accuracy metric were evaluated across strategy-dataset combinations using Spearman correlation analysis.

To assess differences among MEMs after accounting for model size, separate linear models were fitted for each accuracy metric and model-size measure:

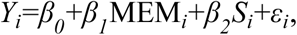

where *Y_i_* denotes the raw accuracy score for strategy–dataset observation *i*, MEM*_i_* denotes the model extraction method and *S_i_* denotes model size. Reaction, gene and metabolite numbers were evaluated separately in independent models.

For each fitted model, size-adjusted MEM scores were predicted at the mean model size across all observations included in the corresponding analysis. Standard errors and corresponding 95% confidence intervals were derived from the fitted linear models. Observed MEM-level scores before adjustment were calculated as the mean raw score across all available strategy-dataset observations. The same observed scores were used as the before-adjustment reference for analyses based on reaction, gene and metabolite numbers. Before- and after-adjustment scores were compared separately for each model-size measure.

## Statistical analysis

Three-factor analysis of variance was used to evaluate the effects of model extraction method (MEM), data preprocessing method (Data) and gene expression threshold (Threshold) on raw accuracy scores. The analysis included the 24 threshold-dependent strategies comprising ftINIT, GIMME and iMAT, two data preprocessing methods and four gene expression thresholds; rFASTCORMICS was excluded because it does not require an externally specified threshold. MEM, Data and Threshold were treated as categorical variables, and a main-effects model without interaction terms was fitted using the MATLAB anovan function with the linear model specification. Analyses were performed separately for individual datasets and for cross-dataset score values. For factors with significant main effects, pairwise comparisons were performed using multcompare, with P<0.05 considered statistically significant.

## Code availability

The code can be accessed at https://github.com/ChenYuGroup/scGEM_benchmark.

