## Supplementary information for "A benchmarking framework for single-cell genome-scale metabolic model construction"

### Supplementary Notes

Supplementary Note 1: Exclusion criteria and settings for benchmarking model extraction methods.

### Supplementary Figures

Supplementary Fig. 1 | Raw accuracy scores grouped by scGEM construction factor using alternative housekeeping gene sets and NMI.
Supplementary Fig. 2 | Factor-level accuracy profiles derived from the 26 construction strategies for single-cell genome-scale metabolic model (scGEM).
Supplementary Fig. 3 | Variance explained by scGEM construction factors across accuracy metrics.
Supplementary Fig. 4 | Expression-perturbation sensitivity assessed using alternative housekeeping gene sets and NMI.
Supplementary Fig. 5 | Pairwise similarity of construction-strategy performance across datasets for housekeeping gene coverage, metabolic task completion and essential gene prediction.
Supplementary Fig. 6 | Dataset-specific marker gene coverage across scGEM construction strategies.
Supplementary Fig. 7 | Dataset-specific effects of scGEM construction factors on clustering performance.
Supplementary Fig. 8 | Variance decomposition of construction-factor effects on NMI-based clustering performance.
Supplementary Fig. 9 | Comparison of NMI-based clustering performance between scGEMs and expression-based clustering.
Supplementary Fig. 10 |Model sizes across scGEM construction strategies.
Supplementary Fig. 11 | Pairwise associations among scGEM model-size measures.
Supplementary Fig. 12 | Associations between model size and raw accuracy metrics across scGEM construction strategies.
Supplementary Fig. 13 | Associations between model size and functional accuracy metrics excluding ftINIT-based strategies.
Supplementary Fig. 14 | MEM performance before and after adjustment for metabolite count.

### Supplementary Notes

#### Supplementary Note 1 | Exclusion criteria and settings for benchmarking model extraction methods

We initially considered 14 candidate model extraction methods (MEMs) for context-specific genome-scale metabolic model reconstruction. Following previously established classifications, these methods were organized into three major families according to their underlying mathematical objectives: GIMME-like, iMAT-like and MBA-like methods[1-3]. We additionally considered thermoKernel-based reconstruction. Methods were assessed according to algorithmic distinctiveness, software availability, compatibility with Human-GEM and single-cell transcriptomic data, computational scalability and suitability for automated reconstruction.

The GIMME-like family comprised GIMME[4], GIMMEp[5] and GIM$^3$E[6]. These methods penalize the use of reactions associated with low gene expression while maintaining a predefined level of metabolic functionality. GIMME was retained because it is a representative and widely used method in this family and was computationally suitable for automated reconstruction of large numbers of cell-specific models. GIMMEp and GIM$^3$E were not evaluated separately because they extend the same general objective-constrained reconstruction principle and would provide limited additional algorithmic diversity relative to GIMME.

The iMAT-like family comprised iMAT[7], INIT[8], tINIT[9] and ftINIT[10]. iMAT seeks to maximize consistency between expression-derived reaction states and predicted reaction activity without requiring predefined metabolic tasks. INIT-related methods use expression-derived reaction weights to guide model extraction, whereas tINIT additionally requires the resulting model to satisfy predefined metabolic tasks. ftINIT is a computationally accelerated implementation of the tINIT workflow. iMAT and ftINIT were both retained because they represent distinct task-independent and task-constrained implementations within the iMAT-like family. INIT and tINIT were not included separately because ftINIT preserves the task-constrained reconstruction principle while providing greater computational efficiency for large-scale single-cell model construction.

The MBA-like family comprised MBA[11], mCADRE[12], FASTCORE[13], FASTCORMICS[14], rFASTCORMICS[15] and CORDA[16]. These methods use predefined or expression-derived reaction-confidence levels to identify context-specific subnetworks while preserving flux consistency. MBA and mCADRE were excluded because their iterative reconstruction procedures were not computationally compatible with the scale of the present benchmark. Previous reports have described substantial computational requirements for MBA-based reconstruction[12], and in our pilot analyses mCADRE required approximately three to four days to reconstruct a single model under the computational conditions used in this study. CORDA was also excluded because pilot reconstructions required approximately four to seven hours per model. These runtimes were considered unsuitable for repeated reconstruction of large numbers of cell-specific models.

FASTCORE was not evaluated separately because it requires a predefined set of core reactions and does not provide a complete workflow for processing RNA-seq expression data. FASTCORMICS extends the FASTCORE principle by deriving reaction-confidence information from discretized transcriptomic data, whereas rFASTCORMICS provides an RNA-seq oriented reconstruction workflow with automated expression discretization and gene mapping. rFASTCORMICS was therefore selected as the representative MBA-like method because it was directly applicable to the input data used in this study and was suitable for repeated automated reconstruction.

We additionally considered thermoKernel as implemented in the XomicsToModel[17] workflow. This approach constructs thermodynamically and flux-consistent context-specific models and showed moderate computational requirements when feasible inputs were available. However, in pilot analyses, some input profiles did not yield feasible models without additional inspection or adjustment of intermediate constraints. Because the benchmark required a standardized and fully automated workflow across large numbers of cell-specific reconstructions, thermoKernel-based reconstruction was not included in the final comparison.

The final benchmark therefore included four MEMs: GIMME, iMAT, ftINIT and rFASTCORMICS. These methods represented distinct objective-constrained, expression-consistency, task-constrained and core-reaction-based reconstruction principles while satisfying the requirements for software accessibility, compatibility with Human1 and computational scalability.

### Supplementary Figures

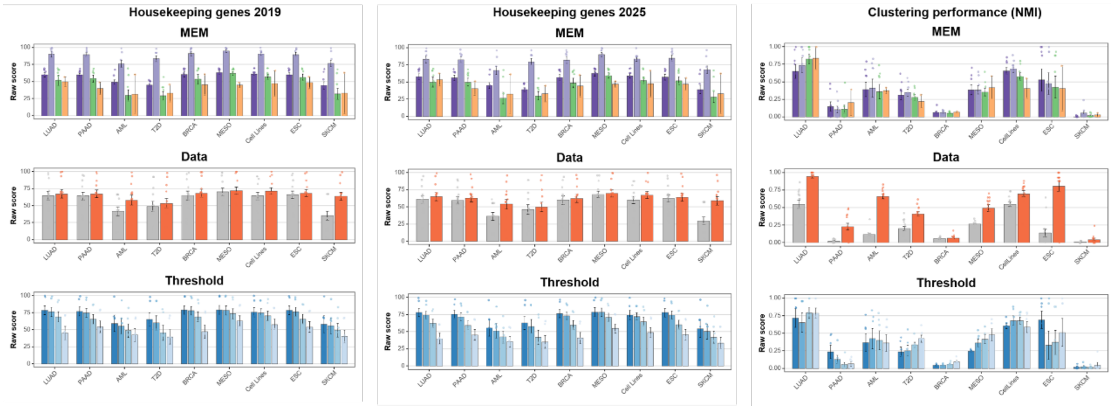


**Supplementary Fig. 1 | Raw accuracy scores grouped by scGEM construction factor using alternative housekeeping gene sets and NMI.** Scores are grouped by model extraction method (MEM), data preprocessing method (Data) and gene expression threshold (Threshold). Housekeeping gene coverage is shown using the housekeeping gene set published in 2019 and 2025, and clustering performance is represented by the normalized mutual information (NMI). Threshold comparisons include ftINIT, GIMME and iMAT only; rFASTCORMICS was excluded because it does not require an externally specified gene expression threshold. Within each dataset, bars show the mean raw score across strategies within each factor level and dataset, error bars indicate the standard error of the mean, and points represent individual strategies.

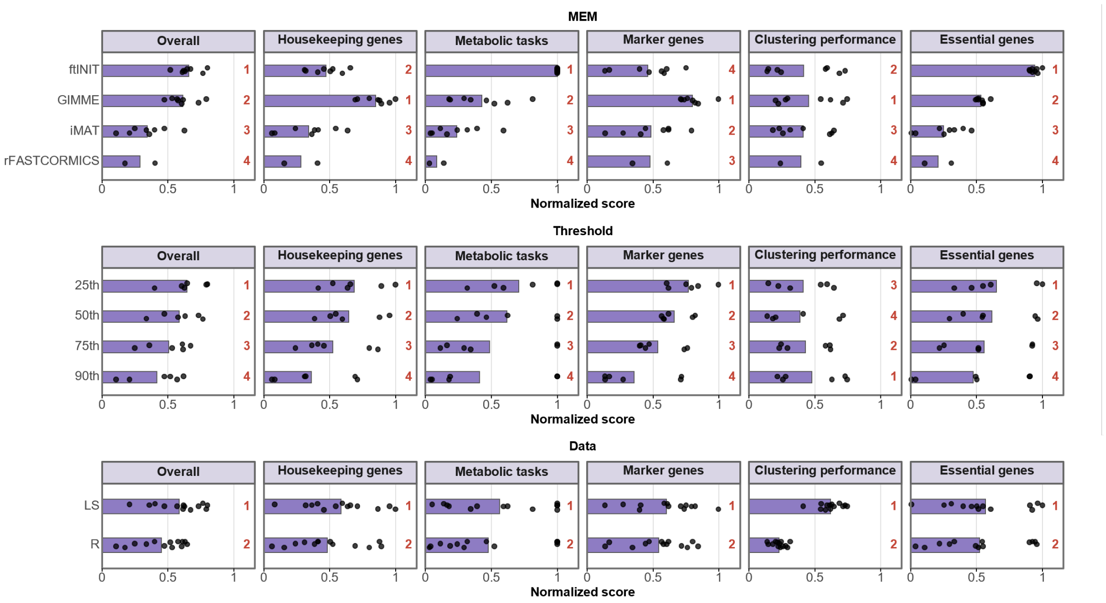


**Supplementary Fig. 2 | Factor-level accuracy profiles derived from the 26 construction strategies for single-cell genome-scale metabolic model (scGEM).** The strategies were grouped by model extraction method (MEM), gene expression threshold (Threshold) and data preprocessing method (Data), and evaluated across five accuracy metrics and the overall score. R denotes the raw expression matrix, whereas LS denotes the expression matrix normalized using Linnorm and imputed using SAVER. Threshold levels correspond to the 25th, 50th, 75th and 90th percentiles of gene expression. For each metric and dataset, raw scores were min–max normalized across strategies and then averaged across datasets to obtain a normalized score for each strategy. Bars show the mean strategy-level score within each factor level, and points represent individual strategies. The overall score was calculated as the unweighted mean of the five metric-level scores. Red numbers indicate the rank of each factor level for the corresponding metric or overall score, with higher scores indicating better performance. The MEM comparison comprised eight strategies for each of ftINIT, GIMME and iMAT and two strategies for rFASTCORMICS. The Threshold comparison comprised six strategies per threshold, whereas the Data comparison comprised 13 strategies per preprocessing method. rFASTCORMICS was excluded from the Threshold comparison because it does not require a predefined gene expression threshold.

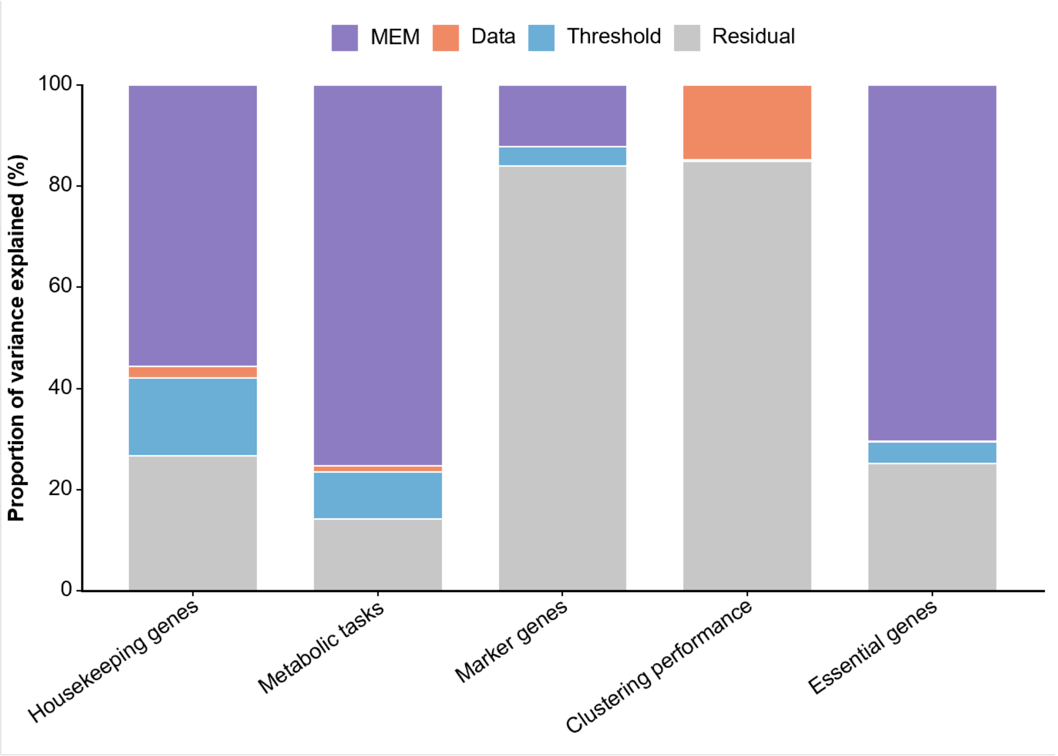


**Supplementary Fig. 3 | Variance explained by scGEM construction factors across accuracy metrics.** Stacked bars show the proportions of total variance associated with model extraction method (MEM), data preprocessing method (Data), gene expression threshold (Threshold) and residual variation across five accuracy metrics. For each metric, dataset-specific scores were pooled across datasets and analyzed using a main-effects linear model. rFASTCORMICS was excluded because it does not require a predefined gene expression threshold. Residual variation includes between-dataset variation, unmodeled factor interactions and other unexplained variation.

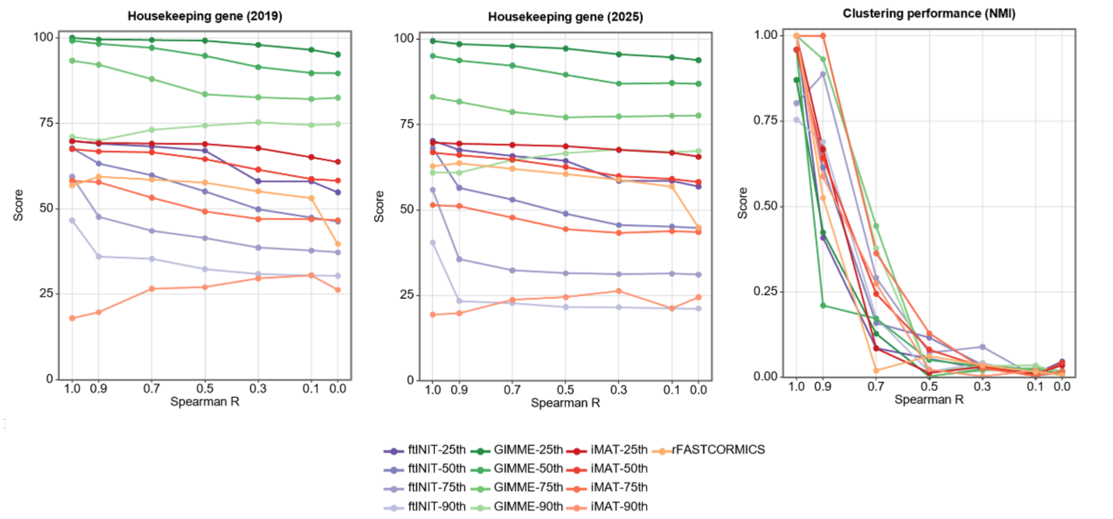


**Supplementary Fig. 4 | Expression-perturbation sensitivity assessed using alternative housekeeping gene sets and NMI.** Sensitivity of LS-based scGEM construction strategies to progressive expression perturbation in the LUAD dataset. LS denotes expression matrices normalized using Linnorm and imputed using SAVER. The unperturbed expression matrix (R = 1) served as the reference, and perturbed matrices were generated at target Spearman rank correlations of $\rho = 0.9$, 0.7, 0.5, 0.3, 0.1 and 0 with the original matrix. Each line represents one construction strategy. Panels show raw housekeeping gene coverage based on the housekeeping gene sets published in 2019 and 2025 and clustering performance quantified using normalized mutual information (NMI).

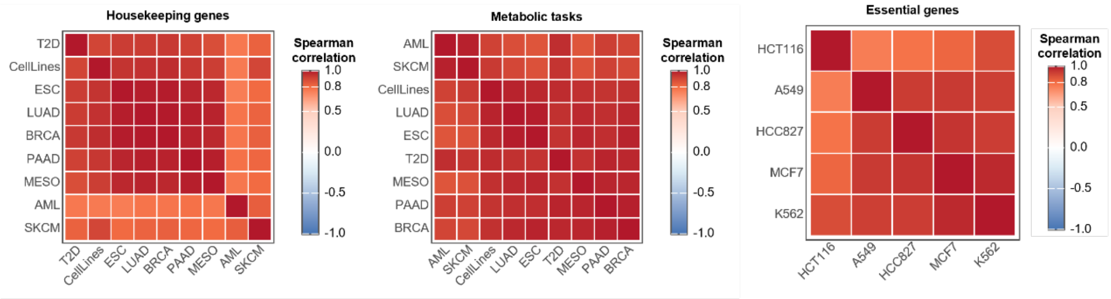


**Supplementary Fig. 5 | Pairwise similarity of construction-strategy performance across datasets for housekeeping gene coverage, metabolic task completion and essential gene prediction.** Each heatmap cell shows the Spearman rank correlation between the scores of the 26 construction strategies for a pair of datasets. The analyses included nine datasets for housekeeping gene coverage and metabolic task completion and five datasets for essential gene prediction.

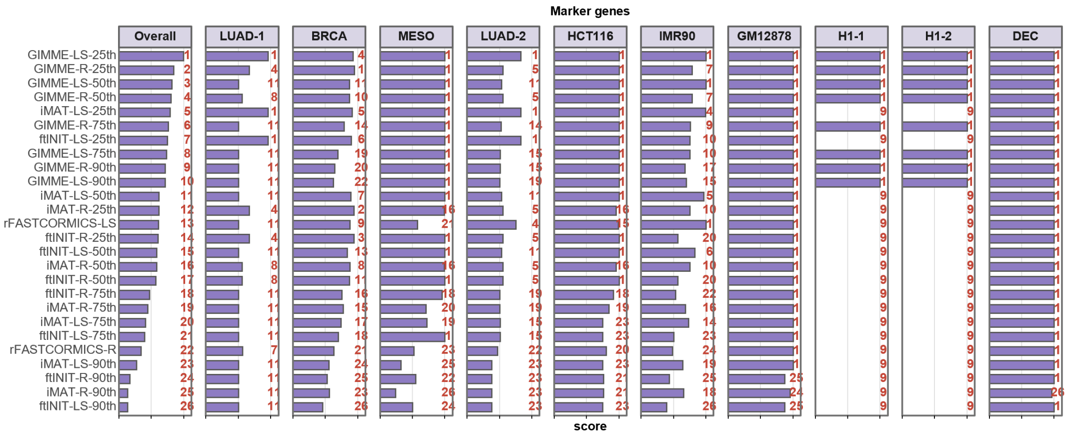


**Supplementary Fig. 6 | Dataset-specific marker gene coverage across scGEM construction strategies.** Horizontal bars indicate the mean proportion of dataset-specific metabolic marker genes retained in the constructed scGEMs. Strategies are ordered by overall performance, and red numbers indicate their rank within each panel. R denotes raw expression data, and LS denotes expression data normalized using Linnorm and imputed using SAVER. LUAD-1 and LUAD-2 correspond to the LUAD dataset and the A549 and H1437 cell lines within the CellLines dataset, respectively, whereas H1-1 and H1-2 correspond to H1 cells within the CellLines and ESC datasets, respectively.

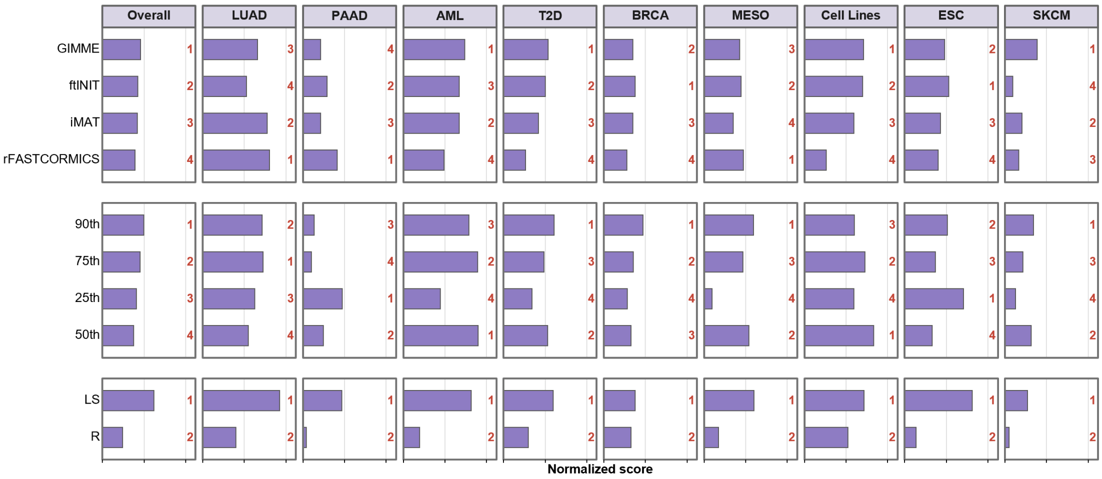


**Supplementary Fig. 7 | Dataset-specific effects of scGEM construction factors on clustering performance.** Normalized clustering scores summarized by model extraction method (MEM; top), gene expression threshold (Threshold; middle) and data preprocessing method (Data; bottom) across all applicable construction strategies. Scores for each factor level were averaged over the other applicable construction factors. The Overall column shows the mean score across the nine datasets. Bars indicate normalized clustering scores, and red numbers indicate ranks within each panel; higher scores indicate better performance. rFASTCORMICS was excluded from the Threshold comparison because it does not require an externally specified gene expression threshold. R denotes raw expression data, and LS denotes expression data normalized using Linnorm and imputed using SAVER.

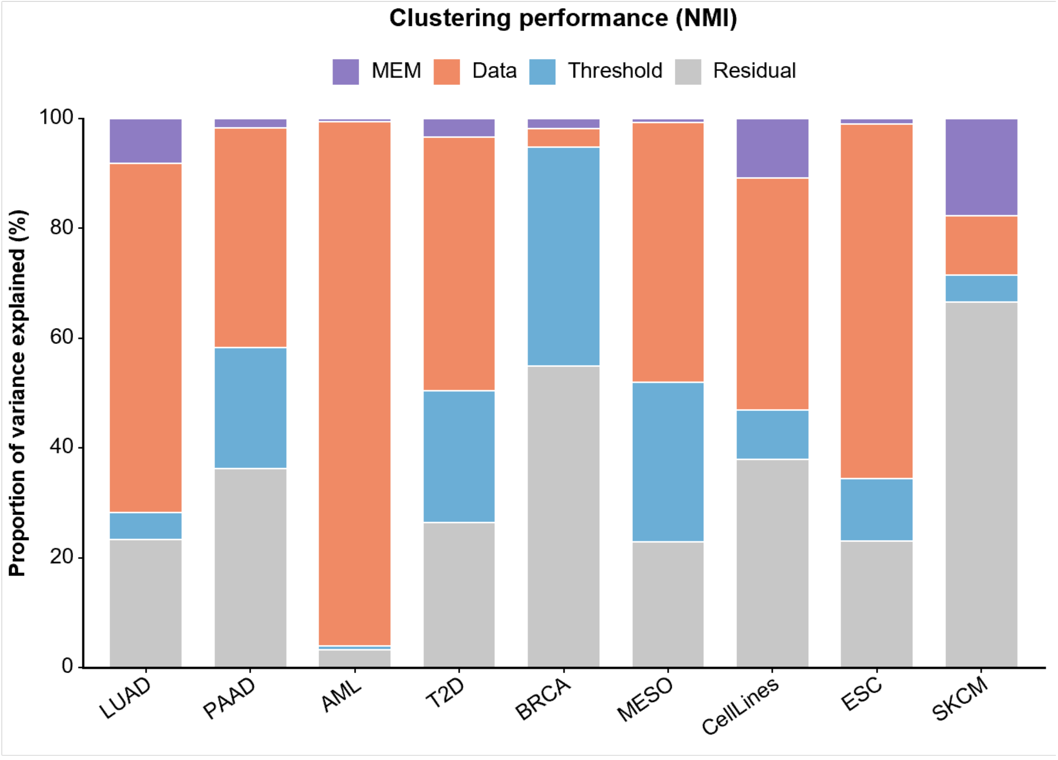


**Supplementary Fig. 8 | Variance decomposition of construction-factor effects on NMI-based clustering performance.** Variance decomposition of raw normalized mutual information (NMI) values across benchmark datasets. Stacked bars show the proportion of variance explained by model extraction method (MEM), data preprocessing method (Data) and gene expression threshold (Threshold), with unexplained variation represented by the residual. rFASTCORMICS was excluded from analyses involving Threshold.

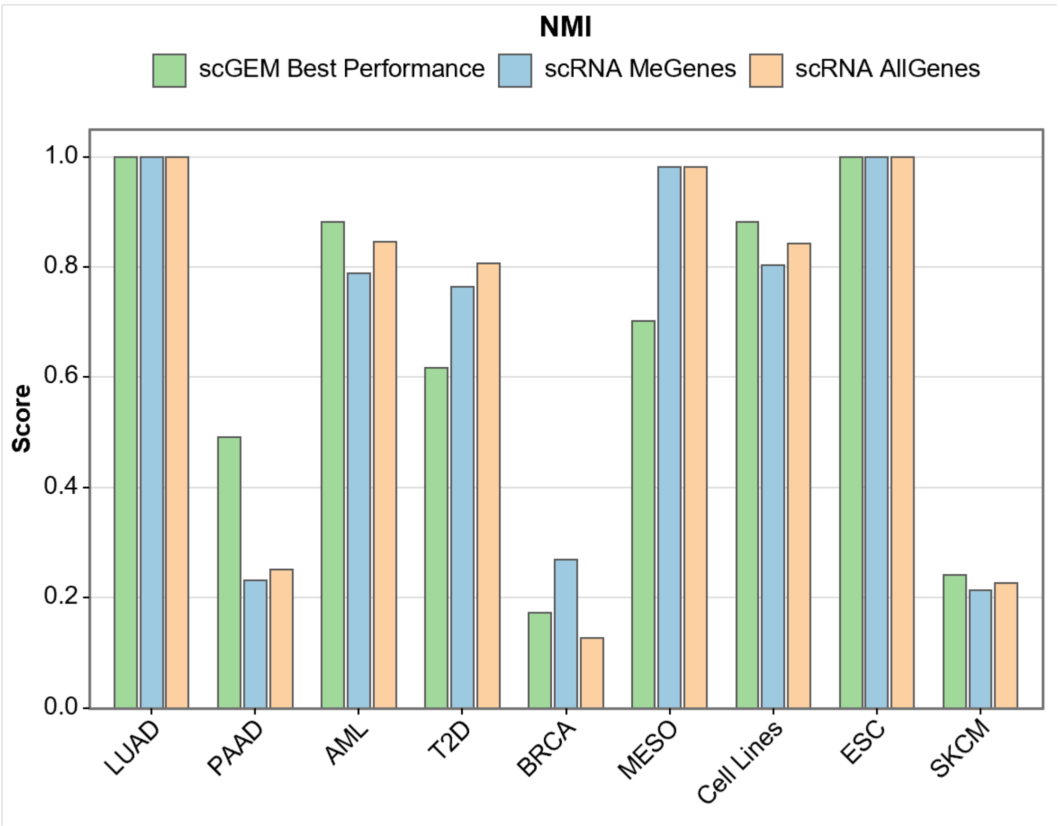


**Supplementary Fig. 9 | Comparison of NMI-based clustering performance between scGEMs and expression-based clustering.** For each dataset, the scGEM result corresponds to the construction strategy that achieved the highest normalized mutual information (NMI) value. Expression-based clustering was performed using scRNA-seq expression profiles restricted to metabolic genes represented in Human-GEM or using all detected genes. Bars show the corresponding raw NMI values across the nine benchmark datasets.

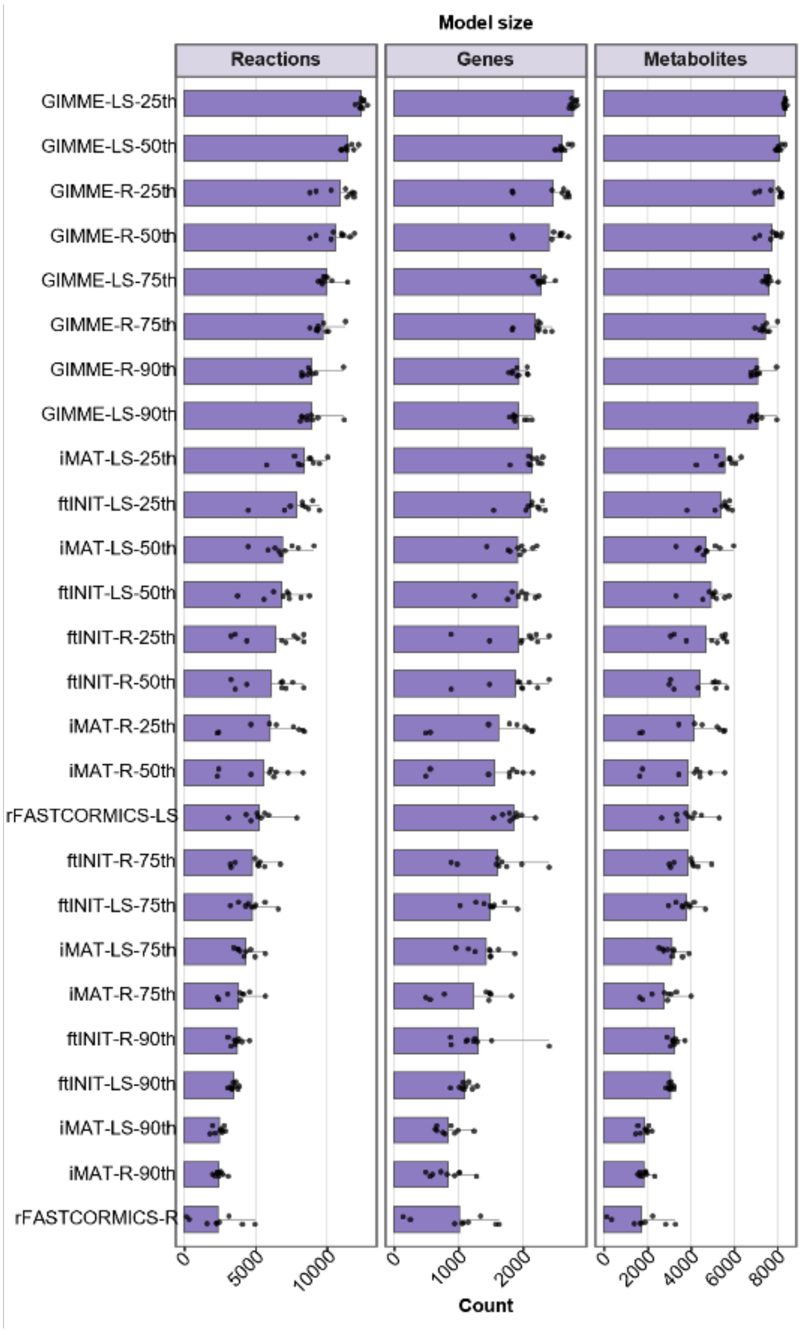


**Supplementary Fig. 10 | Model sizes across scGEM construction strategies.** Numbers of reactions, genes and metabolites in scGEMs generated using the 26 construction strategies. For each strategy and dataset, model-size measures were averaged across cell-specific models. Bars show the mean dataset-level value across benchmark datasets, and points show values for individual datasets. Strategies are ordered according to their mean reaction number, with the same order retained across panels. R denotes the raw expression matrix, and LS denotes the matrix normalized using Linnorm and imputed using SAVER.

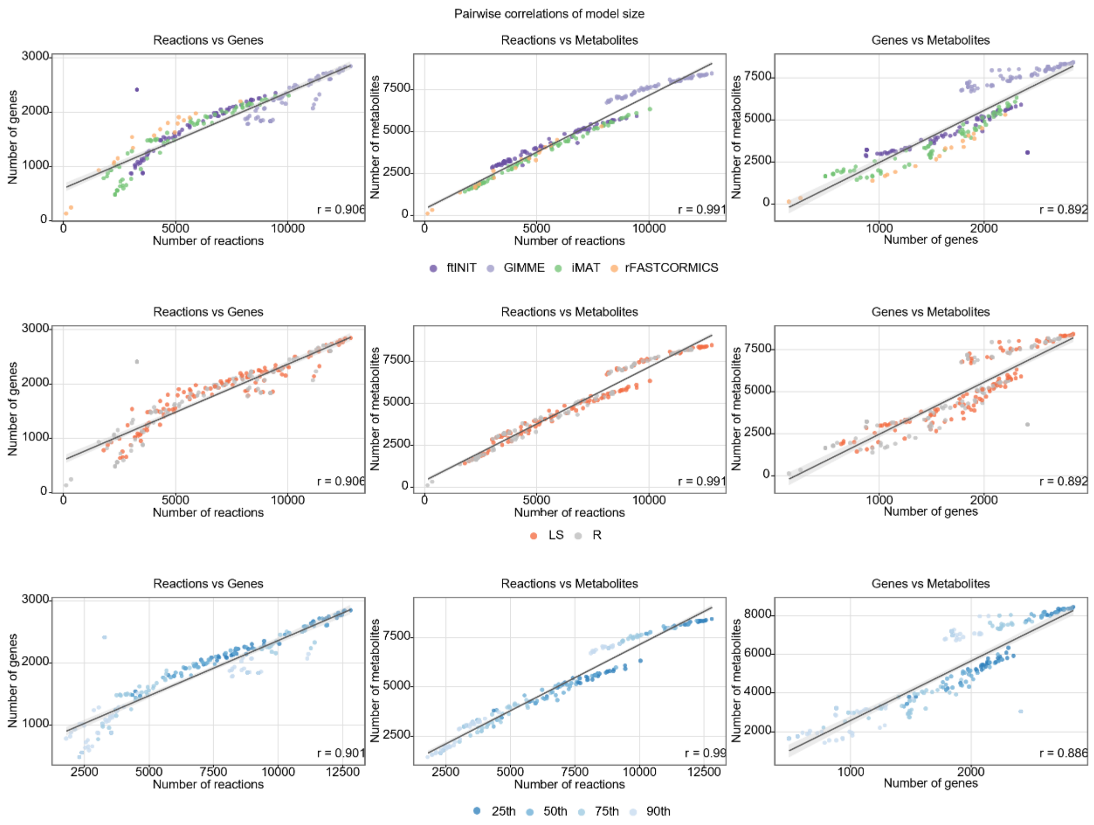


**Supplementary Fig. 11 | Pairwise associations among scGEM model-size measures.** Pairwise associations among dataset-level mean reaction, gene and metabolite counts for scGEMs. For each construction strategy and dataset, model-size measures were first averaged across cell-specific models, and each point represents one resulting strategy–dataset observation. Columns compare reaction and gene counts, reaction and metabolite counts, and gene and metabolite counts. Rows show the same comparisons with points colored by model extraction method (MEM), data preprocessing method (Data) or gene expression threshold (Threshold). rFASTCORMICS was excluded from the Threshold comparisons because it does not require an externally specified threshold. Grey lines show linear regression fits, with shaded areas indicating 95% confidence intervals. Spearman correlation coefficients (r) are reported in the corresponding panels.

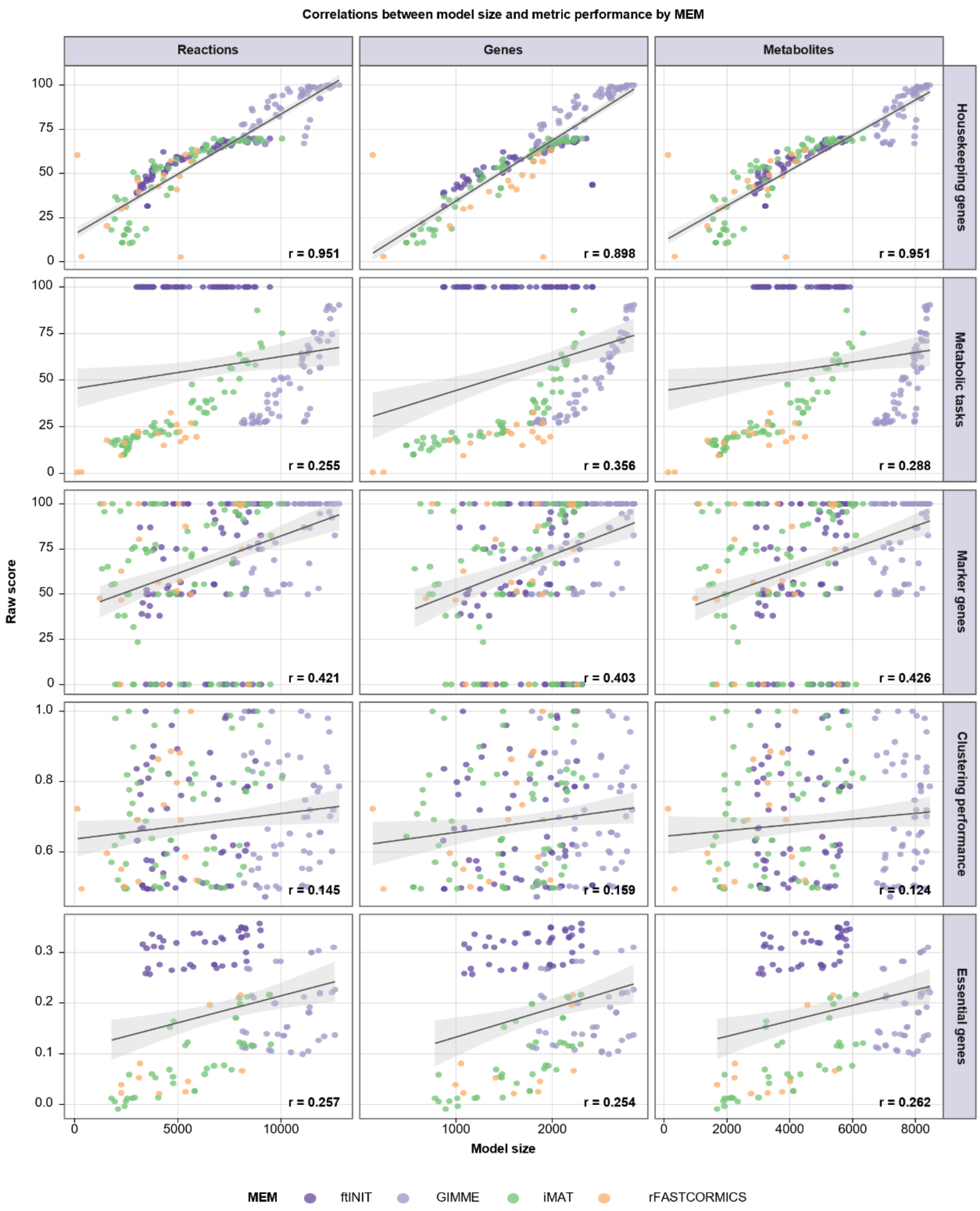


**Supplementary Fig. 12 | Associations between model size and raw accuracy metrics across scGEM construction strategies.** Associations of dataset-level mean reaction, gene and metabolite counts with raw accuracy metrics across strategy-dataset combinations. For each construction strategy and dataset, model-size measures were averaged across cell-specific models, and each point represents the resulting dataset-level model-size and accuracy values. Points are colored by model extraction method (MEM). Grey lines show linear regression fits, with shaded areas indicating 95% confidence intervals. Spearman correlation coefficients (r) are reported in the corresponding panels. Housekeeping gene coverage is shown using the housekeeping gene set published in 2022, and clustering performance is represented by the adjusted Rand index (ARI).

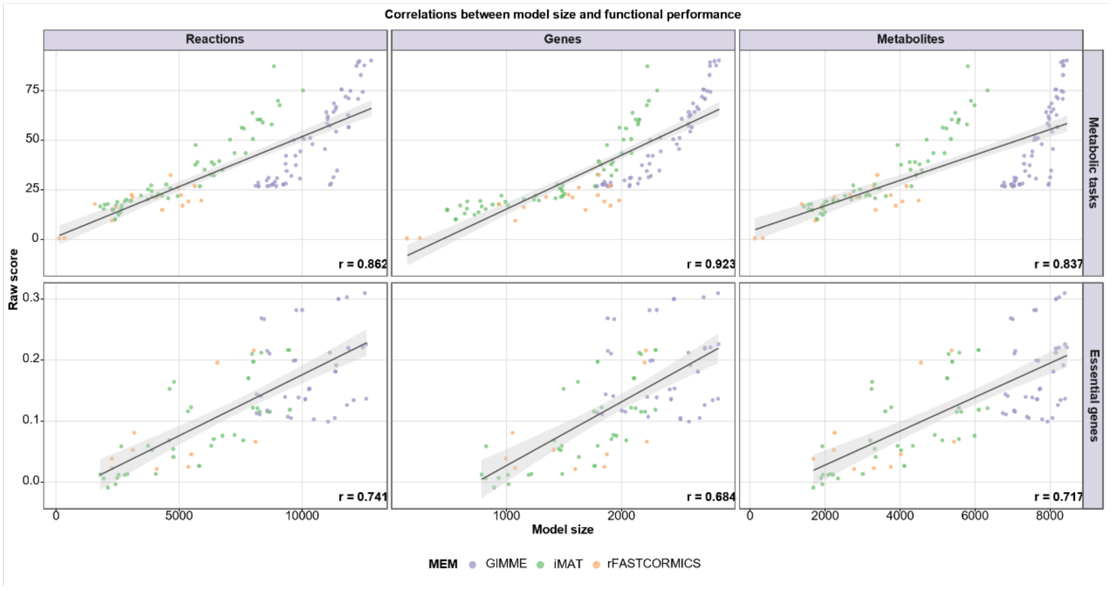


**Supplementary Fig. 13 | Associations between model size and functional accuracy metrics excluding ftINIT-based strategies.** Associations of dataset-level mean reaction, gene and metabolite counts with raw metabolic task completion and essential gene prediction scores after excluding ftINIT-based strategies. Rows correspond to the two functional accuracy metrics, and columns correspond to the three model-size measures. For each remaining construction strategy and dataset, model-size measures were averaged across cell-specific models, and each point represents the resulting strategy–dataset observation. Points are colored by model extraction method (MEM). Grey lines show linear regression fits, with shaded areas indicating 95% confidence intervals. Spearman correlation coefficients (r) are reported in the corresponding panels. Only datasets for which the reference data required to calculate the corresponding accuracy metric were available were included.

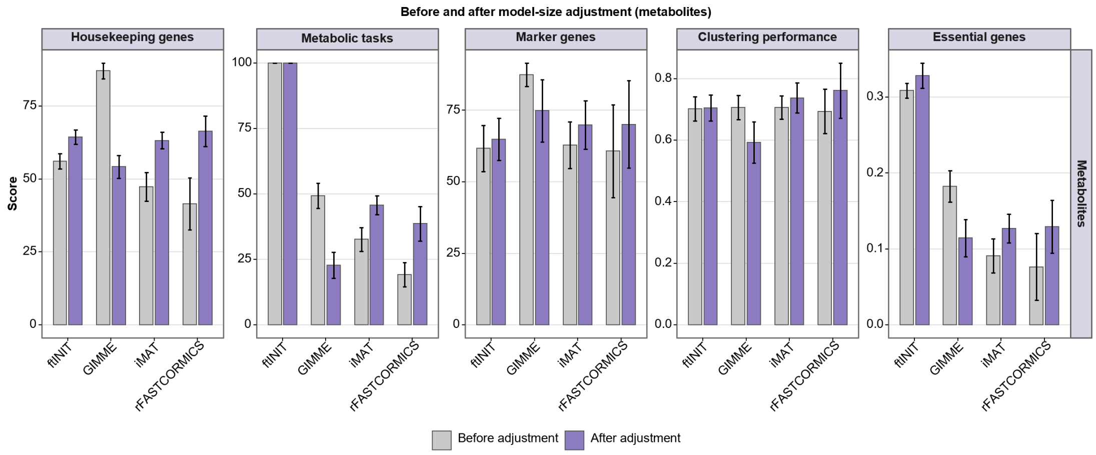


**Supplementary Fig. 14 | MEM performance before and after adjustment for metabolite count.** Separate linear models were fitted for each accuracy metric using the raw metric score as the response and model extraction method (MEM) and metabolite number as explanatory variables. Grey bars show unadjusted mean raw scores, and purple bars show adjusted MEM-level scores evaluated at the mean metabolite number of the observations included in each analysis. Error bars indicate 95% confidence intervals.
